# Genome-wide transcriptional silencing and mRNA stabilization allow the coordinated expression of the meiotic program in mice

**DOI:** 10.1101/2024.02.08.579523

**Authors:** Laura Bellutti, Edith Chan Sock Pen, Marie-Justine Guerquin, Sébastien Messiaen, Elena Llano, Victoria Cluzet, Ihsan Dereli, Antoine Rolland, Emmanuelle Martini, Attila Tóth, Alberto M. Pendás, Frederic Chalmel, Gabriel Livera

**Author notes:** Joint Authors (Laura Bellutti, Edith Chan Sock Pen). Laura Bellutti, Université Paris Cité, CNRS, Institut Jacques Monod, F-75013 Paris, France.

## Abstract

The transcriptional dynamic of mammalian cells when these transit from the ubiquitous mitotic to a meiotic specific program is key to understand this switch central to sexual reproduction. By quantifying active RNA polymerase II and nascent transcripts using single cell dataset and ethynyl-uridine pool-down with sorted cells from synchronised testes, we detailed the transcriptional activity of murine male germ cells. When spermatogonia differentiate, transcription slows down, reaching minimal activity at meiotic entry and resumes during pachytene stage. This event, we termed EMLT (for early meiotic low transcription), is distinct from the silencing of sex chromosomes as it is independent of Setdb1 though it is accompanied by the same chromatin mark, H3K9me3. EMLT is delayed in Stra8KO but occurs in mutants altering meiotic chromosome structure or DSB formation or repair. By comparing transcript abundance and neotranscription we unveil a massive event of mRNA stabilization that parallels EMLT. Altogether our data indicate that meiosis is initiated with a nearly silent genome and we propose that the stabilization of transcripts at that time facilitates the meiotic entry by synchronising the expression of several meiotic sub-programs.

## INTRODUCTION

Spermatogenesis relies on specific and complex transcriptional programs to orchestrate the production of male gametes (1). It can be divided into a mitotic phase, a meiotic phase and a post-meiotic phase. Spermatogonial stem cells multiply and enter a differentiation pathway. In the mouse, this leads to a strictly regulated differentiation of type A spermatogonia into type A1-A4, intermediate and B spermatogonia, through successive cell divisions (2). Type B spermatogonia are the precursors of preleptotene cells entering the premeiotic S phase. The spermatocytes undergo two consecutive meiotic divisions allowing the production of haploid spermatids (3). Finally, the so-called “spermiogenesis” includes several morphological transformations of spermatids generating the highly differentiated spermatozoa (4). Each step of spermatogenesis requires a regulated and dynamic expression of specific transcripts, which contributes to the high diversity of testicular transcriptome (5–7). Additionally, several transcriptional arrests and post-transcriptional regulations complicate the testicular transcriptome (8). Two transcriptional arrests have been exquisitely characterized: the silencing of sex chromosomes during meiosis, a process termed Meiotic Sex Chromosome Inactivation (MSCI) and a global silencing in post-meiotic elongated spermatids undergoing a tight packaging of the genetic material (9).

A critical step of spermatogenesis consists in the conversion of mitotic spermatogonia into meiotic spermatocytes. Stimulated from Retinoic Acid 8 (STRA8) is a transcription factor mandatory to orchestrate the mitosis-to-meiosis transition (10). Meiosis begins with a long prophase I that is subdivided into different stages: leptotene, zygotene, pachytene and diplotene. During prophase I, programmed DNA double strand breaks (DSB) are generated at specific regions called “hotspots” (11, 12). DSBs are next repaired by meiotic homologous recombination (HR). In parallel, homologous chromosomes align, pair and synapse thanks to the formation of a proteinaceous structure called the synaptonemal complex (SC). At the end of pachytene stage, DSBs are repaired and autosomes are fully synapsed. In contrast, X and Y chromosomes remain largely unsynapsed because of their lack of homology. They are compartmentalised in a heterochromatic structure called sex body. The sex body is transcriptionally inactive due to DNA damage response and epigenetic regulations that induces the MSCI (9, 13–16).

DSBs generation, HR and SC formation are highly regulated and interdependent processes. The expression of meiotic-specific genes allows fine regulation of these events. Indeed, meiotic initiation is characterised by a large genome reprogramming, to switch from a mitotic program to a meiotic one. This is accompanied by elaborated post-transcriptional mechanisms changing mRNA stability and alternative splicing events (8, 17). The mechanisms governing such considerable changes in the transcriptome are yet poorly understood and may be connected to the major events occurring in the meiotic genome. Interestingly, spermatocytes at first stages of prophase I have been reported with a very low or near-extinct transcriptional activity (18, 19). This genome-wide low transcription level, together with the existence of specific meiotic transcripts and transcriptional regulations, is puzzling. Currently, very little is known about this global transcriptional reduction, contrarily to MSCI which is largely characterised. Few transcriptome analyses focused on spermatocytes initiating meiotic prophase I, due to the difficulty in isolating these rare testicular cell populations. Moreover, transcriptome analyses poorly reflect transcription as these are in part blurred by post-transcriptional mechanisms.

Here, we characterised the global reduction of transcriptional activity in the genome of early meiotic cells. We demonstrated that this decrease of transcription precedes and is independent of the key events of prophase I, DSBs and chromosome synapsis, but depends on the meiotic gatekeeper *Stra8*. Interestingly, this low transcriptional activity is accompanied by a broad stabilization of mRNA at the time of meiotic engagement.

## MATERIAL AND METHODS

### Mice and animals

All animal studies were conducted in accordance with the guidelines for the care and use of laboratory animals of the French Ministry of Agriculture (France, APAFIS#32392-2021071211498973). Mice were housed in controlled photoperiod conditions (lights on from 08:00 to 20:00) and supplied with commercial food and tap water ad libitum. Embryos were collected from dated matings. Males were caged with females overnight, and the presence of a vaginal plug was examined the following morning. The following midday was defined as 0.5 days post-conception (dpc). Mice were sacrificed by cervical dislocation and fetuses were removed from uterine horns before gonad isolation under a binocular microscope. Synchronisation of spermatogenesis in NMRI mice was performed as previously described (20). Two mg of 5-ethynyl uridine (EU, Click-iT™ Nascent RNA Capture Kit, Invitrogen) per mouse were injected at 24 days post-partum (dpp) to label nascent RNA. Three hours later mice were sacrificed and cells were FACS-sorted based on their ploidy to isolate spermatogonia, leptotene and pachytene spermatocytes(21, 22). Cells from 2 to 3 animals were pooled together for RNA extraction and capture of nascent RNA.

All transgenic mice used have been previously described (13, 17, 22–28). Gonads from dogs, cats and goats were obtained following castrations performed in veterinary clinics.

### Preparation of spermatocyte chromosome spreads

Testes used for spermatocyte spread preparations were collected from adult mice (2 to 6 months). For wild-type (WT), *Spo11* ^-/-^, *Meiob*^-/-^, *Setdb1* ^-/-^ and *Atr*^fl/-^; *Atm*^-/-^ mice, spermatocyte spreads were prepared as follows. Spermatocytes were manually liberated from tubules (flat razor blades) and resuspended in 0, 1M sucrose. Ovaries used for chromosome spreads were collected at 14.5 dpc, 18.5 dpc embryos and 0 dpp pups. Ovaries were dilacerated in 0, 1M sucrose on slides using needles. Oocytes or spermatocytes were then distributed on slides in humid chambers and fixed with 1% PFA (in H2O, pH 9.2) and 0.1% Triton for 1h30. After fixation, slides were rinsed with H2O 0.4% Photo-Flo 200 (Kodak).

### Immunofluorescence on meiocyte chromosome spreads

After washing chromosome spreads from spermatocytes or oocytes, slides were air-dried before blocking. Slides were incubated for 1h at room temperature with blocking solution (0.2% BSA, 0.2% Gelatin, 0.05% Tween in PBS) and overnight with primary antibodies in blocking solution at room temperature (see Table S1 for antibodies). Secondary antibodies were incubated for 1h30 at 37°C. Slides were stained with DAPI and mounted with Prolongold medium. Imaging was performed using a Leica DM5500 B epifluorescence microscope (Leica Microsystems) equipped with a CoolSNAP HQ2camera (Photometrics) and Leica MMAF software (Metamorph, Molecular Devices). Images were processed with Image J software.

### Immunohistostaining

Gonads were fixed in 4% PFA. The fixed gonads were dehydrated, embedded in paraffin and cut into 5-μm-thick sections. Sections were mounted on slides, dewaxed, rehydrated and boiled for 20□min in citrate buffer (pH 6) prior to immunostaining. Immunofluorescence staining was performed as follows. Sections were blocked for 1h in Gelatin blocking solution (0.2% BSA, 0.2% Gelatin, 0.05% Tween in PBS) and incubated 1h30 at 37°C with primary antibody in blocking solution (see Table S1 for antibodies). After 3 washes in PBS 0.05% Tween, slides were incubated with secondary antibodies, 1h at 37°C. Slides were treated with DAPI and mounted in Prolongold. Imaging was performed using a spinning-disk confocal microscope (Nikon Ti2, CSU-W1) and Metamorph. Images were processed with Image J software.

### RNA Polymerase II quantification

Quantification of RNA Polymerase II, total (POL2) or phosphorylated (pPOL2), on chromosome spreads was performed with ImageJ software. DAPI staining was used to delimit nuclei of cells and the mean intensity of POL2 was quantified. Background was subtracted from the nuclear signal.

Quantification of POL2 and pPOL2 on testes sections was performed with ImageJ software (Plot profile). DAPI staining was used to draw a line crossing the nucleus of the cell. For each section, the intensity level was normalized to background intensity. For POL2 and pPOL2 comparisons, quantification was performed on the same cells.

### RNA-sequencing data analysis

For EU-RNA-seq, capture of nascent RNA was performed according to the manufacturer instruction (Click-iT™ Nascent RNA Capture Kit, Invitrogen). RNA-seq libraries were generated using Next Ultra II Directional RNA Library Prep kit (NEB) and ribosomal RNA depletion according to the manufacturer’s protocols. EU-RNA-seq libraries were made like the RNA-seq but without any depletion. Libraries were sequenced on an Illumina HiSeq2500 aiming at an average of 60 million 100-bp reads per sample. Raw reads from the EU-RNA-seq and RNA-seq datasets were mapped onto the mouse genome (mm10) by using STAR (v2.5.2a). The *StringTie* tool was used for transcript assembly while Ballgown was used for quantification to get a FPKM expression matrix. After quantification, the resulting FPKM expression matrix from the EU-RNA-seq dataset was corrected according to the expression of 5S ribosomal genes which are considered transcriptionally stable during spermatogenesis. The expression matrix was divided by the weighted sum of all 5S ribosomal genes.

### Single-cell RNA-sequencing data analysis

#### Pre-processing and quality control

Mouse scRNA-seq data were downloaded from the EBI (E-MTAB-6946) (29). These included ten samples from mice aged 5 to 42 dpp. Raw sequencing reads were processed and mapped to the mouse (mm10) genomes using *CellRanger* (v4) with default parameters, and all samples were aggregated with cellranger aggr. Outlier cells were removed from the aggregated matrices with *Scater* (30). In parallel, *DoubletFinder* (31) was used for doublet detection and removal on each individual sample. Finally, *Scater* and *Scran* (32) were used to exclude genes detected in less than 10 cells.

The *Seurat* (v3) analysis pipeline (33) was then applied. Data normalization was performed with the LogNormalize function with default parameters, while the 2000 most variable genes were identified with FindVariableFeatures. A linear dimensionality reduction was then performed with RunPCA and a JackStraw procedure was used to identify the number of dimensions to be considered. The FindNeighbors and FindClusters functions were applied to determine cell clusters. Correction of batch effects was performed using the Seurat’s CCA method. Data visualization was finally achieved using the uniform manifold approximation projection (UMAP) dimensional reduction technique applied to the batch effect corrected data. Following identification of main cell populations based on known marker genes, germ cell clusters were subsetted and submitted using the same pipeline as described above, i.e. applying FindVariableFeatures, RunPCA, JackStraw, FindNeighbors, FindClusters and CellCycleScoring. Again, the clusters were annotated based on known marker genes and the resulting annotation was validated by comparison to that of Ernst and collaborators using the hypergeometric statistical test as well as by monitoring germ cell evolution according to sample ages and cell cycle phase.

#### Quantification of neo-transcription and correction of ploidy effect

Next the velocyto tool was used (34), to generate unspliced and spliced count data starting from bam files from Cell Ranger, to quantify neo-transcription and transcription, respectively. Unspliced scRNA-seq data were used as a proxy for neo-transcription measurement in order to study transcriptional activity during spermatogenesis. The number of unspliced counts for each cell was calculated independently for autosomes and sex chromosomes. Since the amount of DNA by cell can directly influence the number of RNA molecules that are produced in each cell, the ploidy of each germ cell subpopulation was taken into account: For undifferentiated and differentiating spermatogonia (2n/1c) and for secondary spermatocytes (1n/2c), the number of unspliced counts were unchanged; for primary spermatocytes (leptotene, zygotene, pachytene, diplotene and metaphase I; 2n/2c) the number of unspliced counts were divided by two; while for spermatids (1n/1c) the number of unspliced counts were multiplied by two.

#### Identification of escaping and silenced genes during EMLT

Since ploidy-corrected unspliced count matrix was employed, intron size may bias the number of quantified reads per gene. To avoid this effect, a cumulative intron size correction was applied. For each gene: i) intron size was obtained by subtracting the cumulative sum of exons using the Bedtools subtractBed function, ii) this value was then used to divide gene count and the ratio was multiplied by an arbitrary constant (1000000). This normalization allows setting of a threshold for expressed gene detection. Genes showing a null variance were discarded. Next, a two-step process was used to identify potential “silenced” and “escaping” genes. Genes were considered “silenced” if their expression values were: i) higher than an expression threshold (median value of the overall count matrix =0.07) in at least one of the relevant stages (Diff Spg 1, Diff Spg 2, ePa 1, ePa 2, lPa and/or Di) and, ii) significantly correlated (one-tailed Pearson correlation test, p-value < 0.05) to the expression profiles of archetypal genes (*Ndufa11*, *Park7*) showing a silenced expression pattern in Le and Zy. Genes were considered “escaping” if their expression values were: i) higher than 0.07 in Le and Zy and ii) significantly correlated (*p-value* < 0.05) to archetypal genes (*Pet2*, *Ccnb3*, *Prdm9*) showing an escaping expression pattern in Le or Zy.

#### RNA stability

RNA stability during meiosis was investigated using both single-cell and bulk RNA-seq approaches. For the scRNA-seq dataset the “spliced” and “unspliced” expression profiles reflect transcription and neo-transcription, respectively, while the “stability” is the ratio of “spliced” to “unspliced” expression levels. In the RNA-seq dataset “RNA-seq” and “EU-RNA-seq” expression profiles reflect transcription and neo-transcription, respectively, while “stability” is the ratio of “RNA-seq” to “EU-RNA-seq” expression levels.

#### Data representation: heatmaps and boxplots

Heatmaps and boxplots were generated using *Ggplot2* (R-package). For heatmaps expression values of each gene were scaled between [0;1] using a 0-max scale (0=0; maximum value = 1). Thanks to this scaling, zero can be regarded as an actual zero. Stability matrix was scaled between [0;1] with the Rescale() function from the *Scales* package.

#### Integration of gene expression with recombination hotspots and epigenetic marks

Double-strand breaks hotspot data as defined by anti-DMC1 ssDNA sequencing data in C57BL/6J mice was downloaded from GEO (GSE75419) (35). The distance of genes to the nearest hotspot out of 19528 recombination sites was computed with the closest function from *BedTools* (36). Two-tailed Wilcoxon tests were used to compare the distance to the nearest hotspot for “silenced” genes or 1000 random intergenic regions. DSB map defined by SPO11-oligonucleotides was also used to compare recombination sites to transcriptional changes (12).

ChIP-seq data for seven histone marks (H3K4me3, H3K4me1, H3K9me2, H3K9me3, H3K27me3, H3K36me3 and H3K27ac) obtained in spermatogenic cells were download from GEO (GSE132446) (37). In this study histone modifications in homogenous synchronous spermatogenic cells of nine or ten spermatogenic stages (undifferentiated spermatogonia, type A1 spermatogonia, type B spermatogonia, mid-preleptotene spermatocytes, leptotene spermatocytes, mid-zygotene spermatocytes, early-pachytene spermatocytes, mid-pachytene spermatocytes, diplotene spermatocytes) were studied. *DeepTools* was used to perform histone modification distribution analysis around “silenced”, “escaping” genes and random intergenic regions. To focus on the events around the TSS, different upstream and downstream distances were used: -2000/+2000 pb for H3K9me3, H3K9me2, H3K27me3 and H3K4me1; -1000/+1000 for H3K27ac and H3K4me3; and 0/+2000 for H3K36me3. The histone modification scores associated with genomic regions of interest (“escaped”, “silenced” genes and random intergenic regions) were generated by *ComputeMatrix* modules. Bins for each cell type per genomic region were summed in the output matrix. Then profiles of histone modification scores for “silenced”, “escaping” genes and random intergenic regions were represented using mean ± sem.

## RESULTS

### Transcriptional quiescence initiates prior meiotic entry

To finely characterize the transcriptional levels across spermatogenesis, we first quantified the active form of RNA Polymerase II, phosphorylated on Serine 2 of its CTD (pPOL2), by immunofluorescence staining (Figure 1A-G). We first measured pPOL2 intensity on chromatin using spermatocyte chromosome spreads (Figure 1A and B). pPOL2 was hardly detected at the beginning of prophase I, at leptotene, and zygotene stages and weakly observed in early pachytene stages. At the beginning of pachynema, pPOL2 was retrieved in few restricted chromosomal domains. Intensity of pPOL2 increased massively during mid-pachytene stage to reach the maximal intensity level at late pachytene and diplotene stages. As expected, pPOL2 remained almost undetectable on X and Y chromosomes during pachytene and diplotene stages (Figure 1A and Supplementary Figure S1). Quantification of pPOL2 intensity in leptotene cells and in the sex body from mid-pachytene cells indicated an equivalent lack of transcriptional activity. This confirms and extends previous observations (9, 19). We decided to name this process of genome-wide pPOL2 reduction, which takes place at the beginning of prophase I, “Early Meiotic Low Transcription” (EMLT), to distinguish it from the MSCI process which concerns exclusively the sex chromosomes that remain silent in later stages (i.e. pachynema and diplonema).

**Figure 1.**
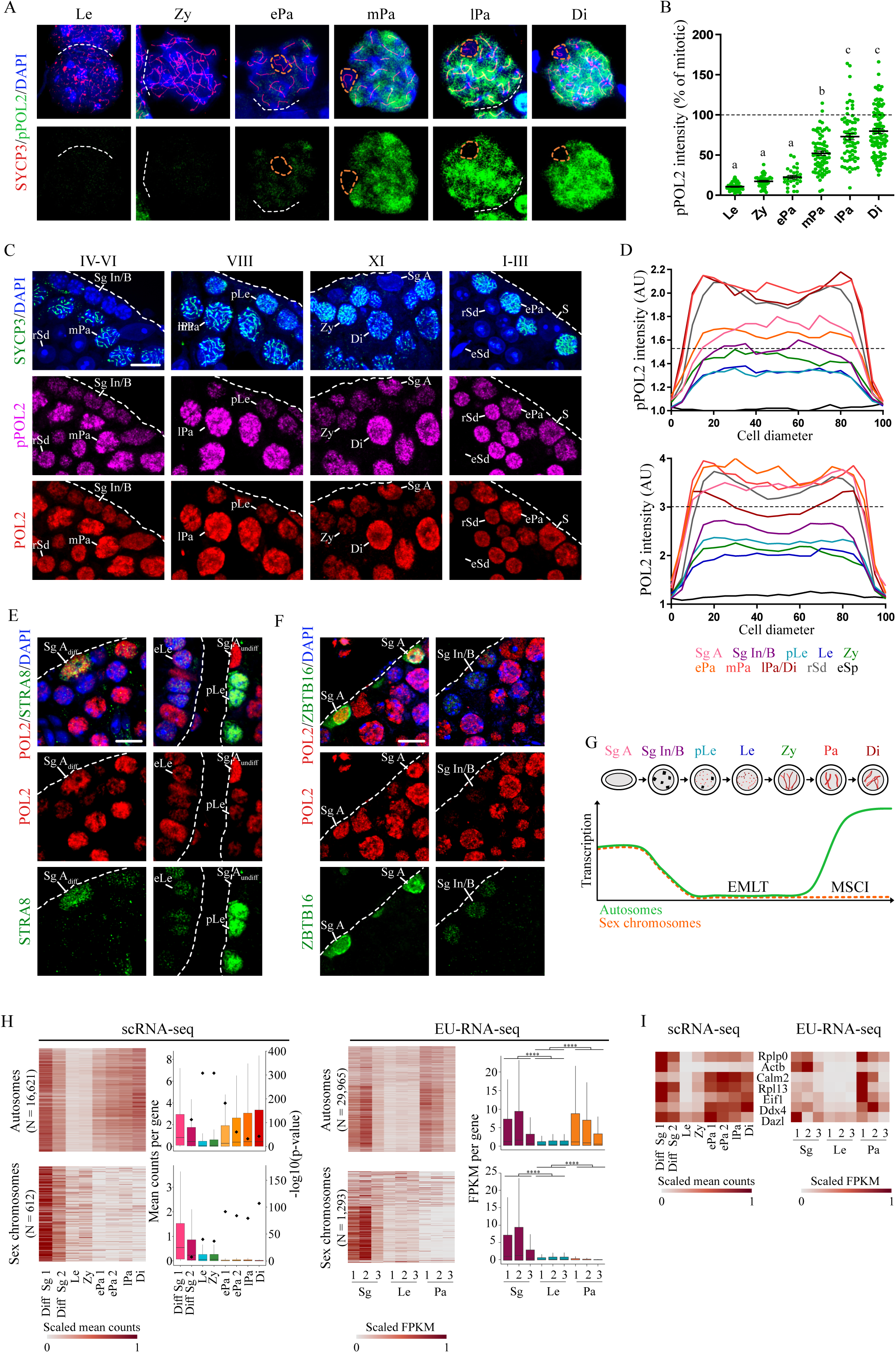
Dynamic of transcription during spermatogenesis. Characterisation of RNA polymerase 2 phosphorylated on Serine 2 (pPOL2) profile during prophase I by co-immunolabelling of pPOL2 (green) and SYCP3 (red) on spermatocytes chromosome spreads (**A**). DNA is stained with DAPI (blue). pPOL2 staining is very low at leptotene (Le), zygotene (Zy) and early-pachytene stages (ePa) and is obvious at mid-pachytene (mPa), late-pachytene (lPa) and diplotene (Di) stages. Sex body (orange dashed circle) is not stained for pPOL2. Quantification of pPOL2 staining (**B**). pPOL2 intensity is normalised to mitotic cells (100%, black dashed line). Each green dot represents one spermatocyte. Spermatocytes were harvested from seven mice; black bars represent means ± sem. For each column, different letters indicate significantly different values (Ordinary one-way ANOVA, multiple comparisons). Immunofluorescence analysis of pPOL2 (magenta) and total RNA polymerase 2 (POL2, red) in testis sections (**C**). Roman numbers indicate the stage of seminiferous epithelium cycle. Prophase I cells are identified thanks to SYCP3 (green) and DNA is stained with DAPI (blue). For A, E and F, dotted white lines delimit seminiferous tubules. Scale bar = 5µm. Quantification of pPOL2 (upper) or POL2 (lower) intensity at each stage of spermatogenesis by line plot analysis in the nuclei (**D**). Each line represents the average of several cells at a specific spermatogenic stage. Sg A= Type A spermatogonia, n=19; Sg Int/B= intermediate and type B spermatogonia, n=26; pLe= preleptotene, n=27; Le=leptotene, n=27; Zy=zygotene, n=22; ePa=early pachytene, n=27; mPa=mid pachytene, n=50; lPa/Di=late pachytene/diplotene, n=32; rSd=round spermatids, n=50; eSd=elongated spermatids, n=4. Black dashed line represents pPOL2 intensity in Sertoli cells (S, n=28). AU=Arbitrary Units. Immunofluorescence analysis of POL2 (red) and STRA8 (green) in testis sections (**E**). DNA is stained with DAPI (blue). Differentiated A spermatogonia (Sg Adiff), pLe and early-leptotene (eLe) cells are stained for STRA8 and show different levels of POL2 intensity. Immunofluorescence analysis of POL2 (red) and ZBTB16 (green) in testis sections (**F**). DNA is stained with DAPI (blue). Sg A and Sp Int/B are stained for ZBTB16 and show different levels of POL2 intensity. Schematic representation of Early Meiotic Low Transcription (EMLT) during spermatogenesis, with a focus on meiotic prophase I (**G**). Transcription level decreases during spermatogonia differentiation to reach minimum at preleptotene and leptotene stages. Transcription is hyperactivated at mid-pachytene and diplotene stages when sex chromosomes remain silent (MSCI). Heatmaps and boxplots of the expression of 17, 233 and 31, 258 genes distributed according to their chromosomal location (autosomes and sex chromosomes) from the unspliced scRNA-seq dataset (average count matrix) and the EU-RNA-seq dataset (FPKM matrix), respectively (**H**). These datasets reflect neo-transcription, and are used to study transcriptional activity during spermatogenesis. In the heatmaps, each row represents a gene, and each column represents a specific spermatogenic cell population (scRNA-seq) or a sample (EU-RNA-seq). For heatmaps, expression values (mean counts or FPKM) of each gene were scaled between [0;1] using a 0-max scale (0=0; maximum value = 1). Boxplots display the mean counts per gene and the FPKM values for unspliced scRNA-seq and for EU-RNA-seq, respectively. One-tailed Wilcoxon tests were used to compare the average counts or FPKM of each spermatogenic stage to the first spermatogenic cells analysed; a second axis representing the -log10(p-value) indicates the test significance for scRNA-seq; **** indicates p < 0.0001 for EU-RNA-seq. Diff Sg = differentiated spermatogonia. Heatmaps representing the gene expression profiles of a subset of emblematic genes robustly expressed in the unspliced scRNA-seq and EU-RNA-seq datasets (**I**).

Next, we exploited the well-characterized seminiferous epithelium cycle to determine when EMLT initiates and ends during spermatogenesis. We thus quantified pPOL2 on testis sections. Type A spermatogonia were clearly stained for pPOL2 (Figure 1C and D and Supplementary Figure S2). In contrast, pPOL2 appeared less abundant in intermediate and type B spermatogonia. Preleptotene and leptotene spermatocytes displayed the lowest pPOL2 staining. During zygonema, the staining increased slightly and progressively to reach at the early pachytene stage an intensity level similar to type A spermatogonia. The maximum intensity of pPOL2 staining was reached at the end of prophase I and in the first round spermatids. This intensity was higher than in A spermatogonia and somatic (Sertoli) cells, As expected, elongated spermatids were completely silenced and devoid of pPOL2 staining due to their high level of DNA compaction. Total RNA Polymerase II (POL2), including non-phosphorylated and phosphorylated forms, had an expression profile similar to that of pPOL2 (Figure 1C and D and Supplementary Figures S1 and S2). Using chromosome spreads, we evidenced that POL2 was also hardly detected on the chromatin during EMLT (Supplementary Figure S1). This indicates that the variations of pPOL2 intensity are due to a reduction in RNA Polymerase II availability and/or loading on chromatin. Interestingly, early pachytene spermatocytes had already reached their maximal intensity for POL2, whereas pPOL2 maximal staining is later, during mid-pachynema (Figure 1D and Supplementary Figure S2B and S3). This suggests a two-step process with first loading POL2 on chromatin and second activation of POL2 (pPOL2) to produce the hyper-transcriptional state observed in late prophase I.

To determine accurately when EMLT was set up during spermatogonia differentiation, we monitored two spermatogonia markers: STRA8 and ZBTB16 (Figure 1E and F). Undifferentiated spermatogonia, that expressed ZBTB16 but not STRA8, and differentiating type A spermatogonia, that expressed ZBTB16 and STRA8 at low level, were stained for POL2. Intermediate and type B spermatogonia, in which ZBTB16 was weakly detected ((38) and personal observations), were faintly stained for POL2. Thus, EMLT initiates during spermatogonia mitotic divisions, more precisely at the transition from type A spermatogonia to intermediate spermatogonia (Figure 1G).

To obtain a direct measure of transcription we analyzed published dataset from single cells (scRNA-seq) and captured nascent transcripts using ethynyl-uridine incorporation and sequencing (EU-RNA-seq). The study of Ernst et al. (2019) did not discard drops containing very few RNAs, termed ‘empty drops’, and was thus well suited to analyze cells with a low transcription (29). Data were downloaded from the EBI database and visualized with UMAP to identify germ cell stages. 34, 078 germ cells were selected and partitioned into 30 clusters (Supplementary Figures S4, S5 and S6). We then analyzed unspliced RNAs as a proxy to monitor neo-transcription. Ploidy effect was corrected as a function of the spermatogenic stage as transcription is strongly associated to ploidy. Gene with null variance were removed and scaled expression of the 17, 233 remaining genes was visualized as a heatmap according to their chromosomal location (autosomes and sex chromosomes). This approach validated the two well-known transcriptional arrests during spermatogenesis, the MSCI during the pachytene stage and the global inactivation during spermiogenesis (Supplementary Figure S6C). Global silencing was observed at the leptotene and zygotene stages (Figure 1H and Supplementary Figure S6C). Total neo-transcription decreased twelve-fold between spermatogonia (Diff Sg1) and leptonema. EU incorporation was performed in post-natal testes with synchronised spermatogenesis and spermatogonia, leptotene and pachytene cells were sorted simultaneously (Supplementary Figure S7A). This approach allowed to minimize contaminations and enrichment of rare stages. Immunostaining for γH2AX and pPOL2 confirmed synchronisation efficiency and that EMLT occurred similarly in synchronised cells (Supplementary Figure S7B). EU-RNA were conjugated to biotin, isolated and sequenced. EU-RNA-seq identified nascent RNA and also evidenced EMLT and MSCI at the leptotene and pachytene stages, respectively (Figure 1G). EU-RNA-seq allowed the analysis of nearly twice as much genes (31, 258) in comparison to scRNA-seq. Comparison of gene transcription with both approaches revealed similar profiles as exemplified by a set of robustly expressed genes (reference genes or germ cell markers, Figure 1I).

Altogether staining for pPOL2 and POL2, pre-mRNA analysis (unspliced scRNA-seq) and nascent RNA analysis (EU-RNA-Seq) confirmed the existence of EMLT. This silencing is initiated prior to meiotic entry and is most pronounced at the leptotene stage in mouse testicular cells.

### EMLT is conserved in mammals and female mouse

As this meiotic genome-wide transcription arrest has been only described in male mice, we sought to determine whether EMLT was conserved among mammalian species. Interestingly, pPOL2 was almost undetectable at the beginning of prophase I in three different mammals: goat, rabbit and dog (Supplementary Figure S8A). As in mice, EMLT was also followed by a hyperactive transcriptional state in pachynema in these mammals. Altogether, this reflects a conserved dynamism of germ cells transcriptional state during spermatogenesis.

Meiosis occurs both during male and female gametogenesis. In female, meiosis is initiated during fetal life and arrests at the diplotene stage around birth. Transcriptional levels during prophase I have been poorly described in females. Thus, we analyzed pPOL2 intensity in oocytes during prophase I (Supplementary Figure S8B and C). We collected embryos at different fetal ages 14.5 and 18.5 days post-conception and at birth and performed chromosome spreads. pPOL2 staining was least abundant in leptotene and zygotene stage oocytes and lower than in somatic cells. pPOL2 staining increased during late prophase I stages. However, there was a large heterogeneity in pPOL2 intensity in female meiocytes whatever the stage. Hence, albeit many oocytes seem to escape a complete transcriptional repression, EMLT appears to be conserved among sexes.

### EMLT mechanism differs from MSCI

In order to search for epigenetic marks correlated to EMLT, we first partitioned genes in two opposite categories based on unspliced RNAs from scRNA-seq. “Silenced” genes exhibited higher transcription levels both before and after EMLT, whereas “escaping” genes were predominantly transcribed specifically during EMLT. Genes were identified by using a correlation-based analysis with archetypal genes, i.e. *Ndufa11* and *Park7* for the “silenced” genes (n=11, 660), and *Pet2*, *Ccnb3* and *Prdm9* for the “escaping” genes (n=79) (Figure 2A and Supplementary Figure S4B). More than 99% of “silenced” genes were located on autosomes (11, 590/11, 660), whereas over 40 % of “escaping” genes (33/79) were found on sex chromosomes, and their profiles were roughly similar when assessed with EU-RNA-seq (Supplementary Figure S9). We next analyzed the profile of several histone marks associated with activation or repression of transcription using published ChIP-seq data (37) (Figure 2B and Supplementary Figure S10). None distinguished the 11, 660 “silenced” from the 79 “escaping” genes. The histone H3K9me2 and me3 modifications around the TSS correlated with the timing of transcriptional quiescence. In particular, H3K9me3 presented a gradual increase from undifferentiated spermatogonia to zygotene stages. This was confirmed by immunostaining in testicular sections (Figure 2C). H3K9me3 staining was observed in patchy areas that likely correspond to pericentromeric heterochromatin. In the rest of the nucleus staining was faint in A spermatogonia and pachytene spermatocytes and most abundant in leptotene and zygotene stages. As H3K9me3 has been proposed to repress transcription and is involved in MSCI, we sought to determine whether the same mechanisms govern MSCI and EMLT. During MSCI, sex chromosomes inactivation is primarily triggered by DNA damage response pathway and the trimethylation of H3K9 by SETDB1 seems to reinforce MSCI (13, 16). We thus analyzed pPOL2 in chromosome spreads from conditional KO obtained with the *Ngn3* -cre for *Setdb1* (*Setb1* ^-/-^) and for both *Atm* and *Atr* (*Atm* ^fl/-^; *Atr*^-/-^). In these animals, MSCI has been reported greatly hampered at pachytene stage. At the leptotene stage, no pPOL2 was detected in the chromatin of both mutant mice indicating that impairment of key factors that govern MSCI do not alter EMLT (Figure 2D). At the early pachytene stage, pPOL2 staining was retrieved in mutants as in wild-type (WT) animals indicating no defect in resuming transcription. To further compare MSCI and EMLT, expression profiles of 612 and 1, 293 genes located on the sex chromosomes were investigated in the unspliced scRNA-seq and the EU-RNA-seq datasets, respectively (Figure 2E). scRNA-seq evidenced that nearly all genes investigated were extinct at the beginning of pachytene stage, many of which had transcriptional depression initiated before meiotic entry. In the EU-RNA-seq, three expression patterns were discernible. The first concerned genes silenced during EMLT but escaping MSCI, the second and predominant cluster was composed of genes silenced during both EMLT and MSCI, and the last cluster consisted of genes escaping EMLT but not MSCI (Figure 2E). Next, we focused on specific genes and gene families on sex chromosomes (Figure 2F). The *androgen receptor* (*Ar*) gene was silenced both during EMLT and MSCI, a profile representative of most genes on sex chromosomes. Nascent RNAs of *Tex16*, *Dusp9* and most genes of the *Mage* family were detected during EMLT, albeit possibly at lower levels than in spermatogonia, but not during MSCI. Genes from the Spin2 family on the contrary were specifically expressed during MSCI. Of note is that Spin2 genes are mono-exonic and were thus discarded from the unspliced scRNA-seq data analysis. Altogether, these data indicate that MSCI is not just a mere extension of EMLT, as some genes on sex chromosomes display different expression dynamics and since key regulators of MSCI do not affect EMLT. Nonetheless repression of transcription may be associated in both cases with H3K9 methylation, possibly using different methylases.

**Figure 2.**
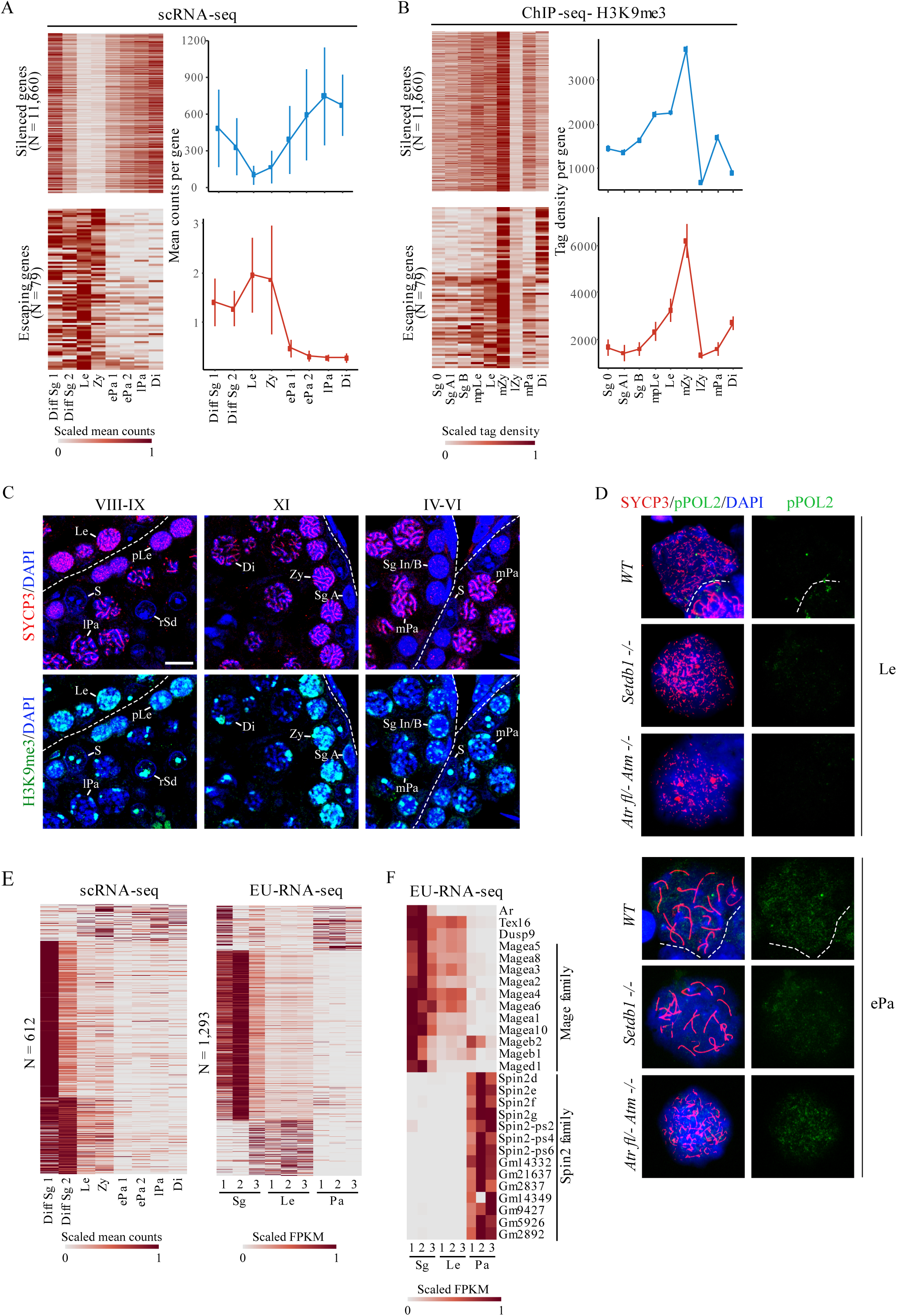
EMLT correlates with H3K9me3 but differs from MSCI. Gene expression profiles of 11, 660 “silenced” and 79 “escaping” genes in the unspliced scRNA-seq dataset represented as heatmaps (scaled mean counts) and profile (mean counts per gene ± sem) (**A**). “Silenced” and “escaping” genes were identified by using a correlation-based analysis with archetypal genes. Quantification of the histone H3K9me3 modification around the TSS of the 11, 660 “silenced” and 79 “escaping” genes according to ChIP-seq data from Chen et al., (2020; GSE132446) (**B**). Heatmaps show the scaled tag density around the TSS from spermatogonia to diplotene spermatocytes: Sg 0 = undifferentiated spermatogonia; Sg A1 = type A1 spermatogonia; Sg B = type B spermatogonia; mpLe = mid-preleptotene spermatocytes; Le = leptotene spermatocytes; mZy = mid-zygotene spermatocytes; lZy = late zygotene spermatocytes; mPa=mid-pachytene spermatocytes; Di = diplotene spermatocytes. The enrichment profiles of “silenced” genes (blue) and “escaping” genes (red) are represented using mean ± sem. Immunostaining for H3K9me3 (green) and SYCP3 (red) in testis sections from adult mice (**C**). DNA is stained with DAPI (blue). Legend is the same as in Figure 1C. Co-labelling with SYCP3 (red) and pPOL2 (green) of spermatocyte chromosome spreads of WT, conditional KO mice for Setdb1 (*Setdb1* -/-) and conditional double KO for *Atr* and *Atm* (*Atr* fl/- *Atm* -/-) (**D**). Conditional inactivation of one *Setb1* and one *Atr* allele was mediated by *Ngn3* -cre (*Ngn3* -cre, *Setdb1* fl/- and *Ngn3* -cre, *Atr* fl/-, *Atm* -/-). All mice show weak pPOL2 staining at leptotene (Le) stage and an increase at early pachytene (ePa) stages. Heatmaps show the scaled expression profiles of genes located on the sex chromosomes in the unspliced scRNA-seq (scaled mean counts) and the EU-RNA-seq (scaled FPKM) datasets according to the scale bars (**E**). Heatmap representing the expression profiles of selected genes of interest for EMLT and for MSCI in the EU-RNA-seq (scaled FPKM) dataset (**F**). This dataset contains mono-exonic genes unlike the unspliced scRNA-seq dataset. The legends of heatmaps in panels A, E and F are the same as for heatmaps in Figure 1H.

### EMLT is independent of chromatin structure and homologous chromosome synapsis

At the beginning of prophase I, spermatocytes are characterized by specific rearrangements of the chromatin structure. Chromosomes form an axe/loop-structure. Sister chromatids are held together by cohesins allowing the formation of axial elements (AE). The establishment of AE begins at early-leptotene stage (39). We thus investigated the relationship between EMLT and chromosome structure using *Rad21L* ^-/-^ and *Stag3* ^-/-^ mice. RAD21L and STAG3 are two important meiotic-specific cohesins (26, 27, 39). Mutation of these cohesins caused an arrest of prophase I progression at or prior to early pachynema. In both *Rad21L* ^-/-^ and *Stag3* ^-/-^, leptotene-like and zygotene-like spermatocytes displayed a low level of pPOL2, similar to that of WT mice (Figure 3A and B). In *Rad21L* mutants, some spermatocytes resembling early pachytene stages presented a more intense staining for pPOL2 (Figure 3B and C), albeit Rad21L mutation renders the staging of these cells uncertain.

**Figure 3.**
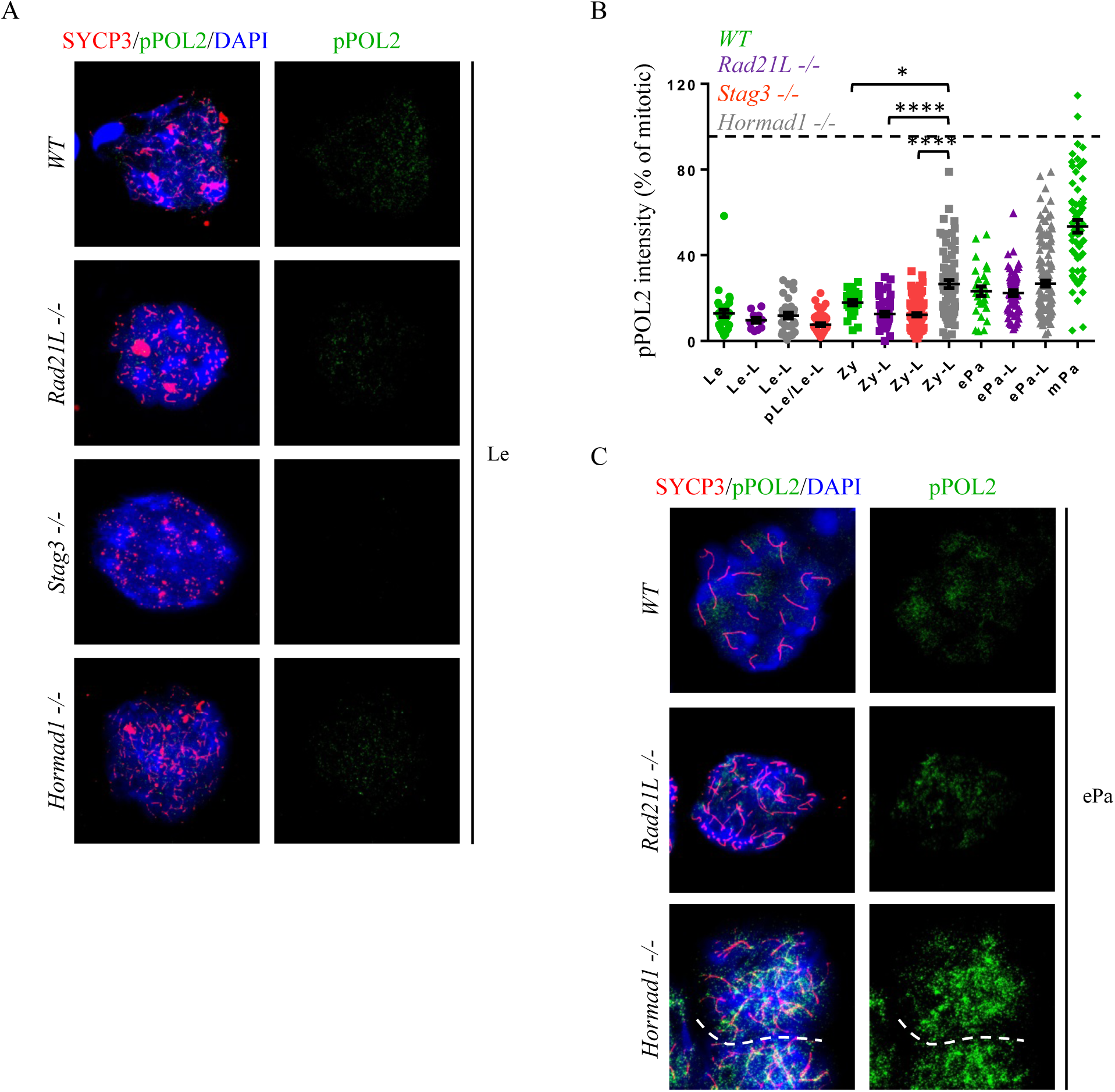
EMLT is independent from chromatin structure and synaptonemal complex. Co-labelling with SYCP3 (red) and pPOL2 (green) on spermatocyte chromosome spreads of WT and KO mice *Rad21L* -/-, *Stag3* -/- and *Hormad1* -/- (**A**). Quantification of pPOL2 intensity evidenced a weak signal at leptotene (Le) or leptotene-like (Le-L) stage (**B**). pPOL2 intensity is normalised to mitotic cells (100%, black dashed line). Black bars represent mean ± sem. WT: n=5; *Hormad1* -/-: n=3; *Stag3* -/-: n=3; *Rad21L* -/-: n=3. *<0, 05, **** P<0.0001 (Unpaired t-test). *In Hormad1* -/-, some early pachytene (ePa) spermatocytes are strongly stained for pPOL2 in comparison to WT and *Rad21L* -/- (**C**).

Proper formation of AEs is mandatory to allow the synapsis of homologous chromosomes through completion of the synaptonemal complex. The HORMA domains protein HORMAD1 localizes to unsynapsed chromosome axes and is removed soon after synapsis (25). The disruption of *Hormad1* caused pairing defects and pachytene arrest. Leptotene and zygotene-like *Hormad1* ^-/-^ spermatocytes were faintly stained for pPOL2, with a quantified intensity similar to WT (Figure 3A and B). Zygotene-like and early pachytene-like mutant spermatocytes appeared more robustly stained for pPOL2 that WT cells (Figure 3C). However, the precise identification of these stages is uncertain in *Hormad1* mutants in which asynapsis signaling is defective. Of note, in WT early pachynema pPOL2 initially reappeared in highly localized areas emanating from the axes of the fully synapsed chromosomes (Supplementary Figure S11). In *Hormad1* ^-/-^ spermatocytes, the reappearance of pPOL2 commenced on both synapsed and unsynapsed axes. This indicates that local synapsis is not a prerequisite for the reinitiation of transcription.

These results indicate that establishment of the meiotic-specific chromatin structure is not at the origin of EMLT. In addition, completion of chromosome synapsis is not essential for concluding EMLT and transcriptional reactivation.

### EMLT is independent of meiotic DSBs

As DSBs can be associated to transcriptional modifications in somatic cells, we speculated that in meiotic cells the formation of hundreds of DSBs might affect transcriptional activity. At the beginning of prophase I, DSBs are generated by the topoisomerase-like transesterase SPO11 at specific sites defined by the methyltransferase PRDM9 (PR/SET domain 9). DSBs are rapidly signaled by phosphorylation of H2AX (γH2AX) by the kinase ATM at leptotene stage. Interestingly, cells stained for γH2AX harbored low pPOL2 (Supplementary Figure S12); this was obvious in early spermatocytes I and in intermediate and B spermatogonia that presented a weak but distinct γH2AX signal as previously reported (40). To investigate if DSBs formation induces or maintains EMLT, we quantified pPOL2 in *Prdm9* and *Spo11* null spermatocytes (Figure 4). In *Prdm9* ^-/-^ mice, DSBs are catalyzed but do not localize correctly (24, 41, 42). In *Spo11* ^-/-^ animals, programmed DSBs are not generated (43, 44). In both cases, mutant spermatocytes arrest at pachytene stage. Quantification of pPOL2 in section of *Prdm9* ^-/-^ testes revealed that EMLT was unchanged as hardly any staining was detected in leptotene and zygotene spermatocytes as in WT cells (Figure 4A and B). This result is in accordance with EMLT being present in dog that lacks *Prdm9* (Supplementary Figure S8A). EMLT was also retrieved similarly in chromosome spreads from *Spo11* ^-/-^ spermatocytes (Figure 4C and D). Of note, in leptotene-like spermatocytes from *Spo11* ^-/-^ mice, pPOL2 intensity is slightly reduced compared to WT (Supplementary Figure S13). This might indicate that in absence of DSB, transcription is even more repressed. Overall, we concluded that EMLT is not induced by programmed DSB.

**Figure 4.**
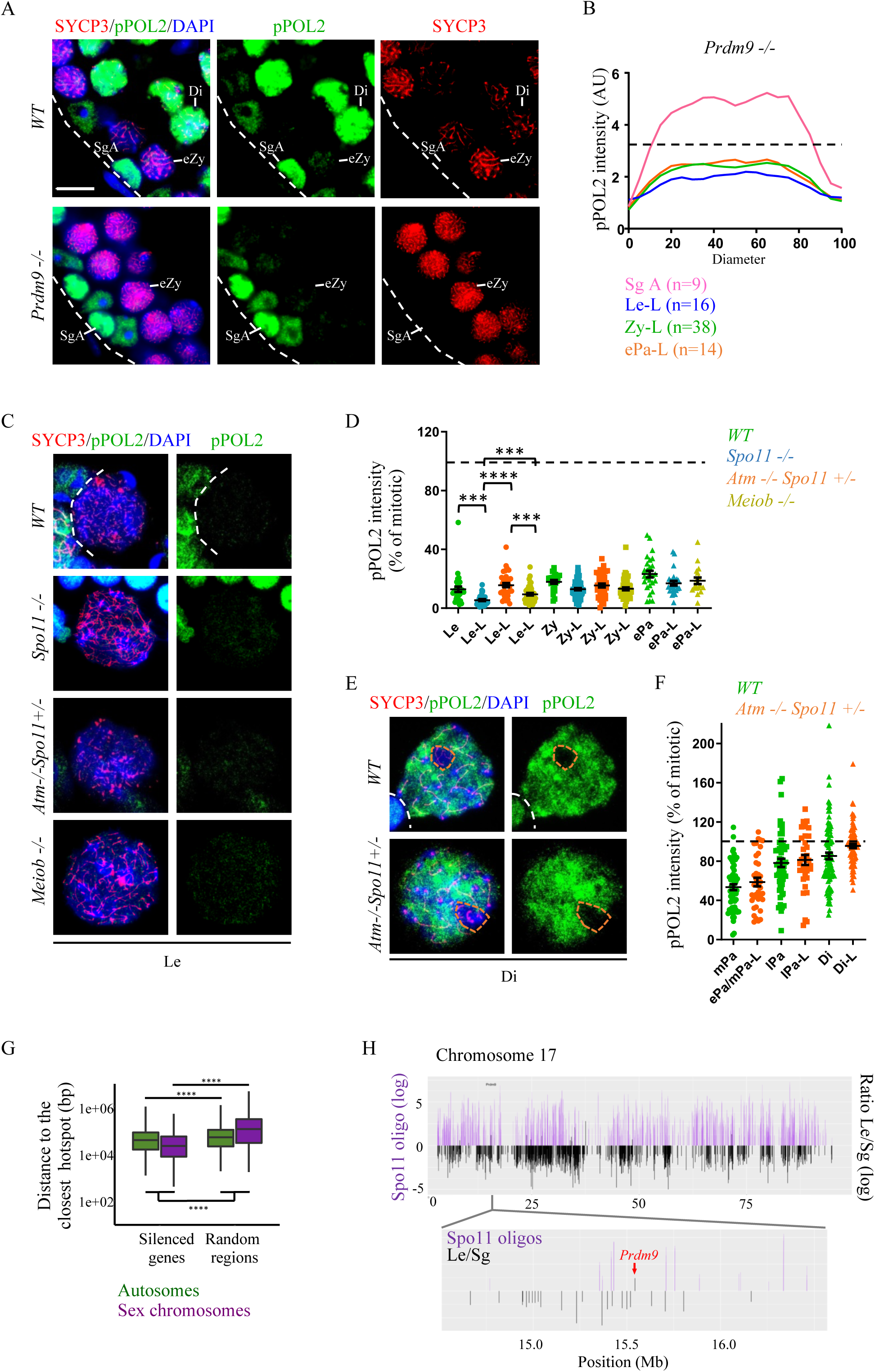
EMLT is independent from Double Strand Breaks. WT and Prdm9 -/- testis sections stained for SYCP3 (red) and pPOL2 (green) (**A**). Prophase I cells are identified thanks to SYCP3 (red) and DNA is stained with DAPI (blue). SgA = spermatogonia A; eZy = early zygotene; Di= diplotene. Dotted white curbs delimit seminiferous tubules. Scale bar= 10µm. Line plot quantification of pPOL2 intensity in Prdm9 null mutants (Prdm9 -/-, n=2 mice) (**B**). For quantification, a line was traced randomly through nucleus and for each point along the line, intensity was quantified using ImageJ software (Plot profile). The horizontal axis represents the diameter of the cells. Each line represents the average of pPOL2 intensity of several cells. Leptotene-like (Le-L, n=16), zygotene-like (Zy-L, n=38) and early-pachytene like (ePa-L, n=14) cells are less stained for pPOL2 than A spermatogonia (Sg A, n=9). AU=Arbitrary Units. Immunofluorescence staining for SYCP3 (red) and pPOL2 (green) of spermatocyte chromosome spreads of WT and KO mice Meiob-/-, Spo11-/- and double transgenic KO mice Atm-/- Spo11+/- (**C**). Quantification of pPOL2 intensity indicates that all mice show weak pPOL2 signal at leptotene (Le) stage (**D**). pPOL2 intensity is normalised to mitotic cells (100%, black dashed line). Black bars represent mean ± sem. WT: n= 5; Meiob -/-: n=3; Spo11 -/-: n=3; Atm-/-Spo11+/-: n=3. ***<0, 001 **** P<0.0001 (Unpaired t-test). Both WT and Atm -/- Spo11 +/- mice show high pPOL2 staining at diplotene stage (Di), except for sex body (orange dotted line)(**E**). Quantification of pPOL2 intensity on spermatocyte chromosome spreads (**F**). Boxplot of the distance (log10) to the closest hotspot (in bp) for the 11, 660 “silenced” genes and 1, 000 random intergenic regions according to their chromosomal location (autosomes in green and sex chromosomes in purple) (**G**). Double-strand breaks hotspot data as defined by anti-DMC1 ssDNA sequencing data in C57BL/6J mice was downloaded from GEO (GSE75419). Two-tailed Wilcoxon tests were used to compare the distance to the nearest hotspot in the different sets of genes. ****: p < 0.0001. Illustration of the distribution of DNA double-strand breaks defined by mapping SPO11 oligonucleotides (Spo11 oligo, purple) and transcriptional changes between leptotene stage and spermatogonia (Ratio Le/Sg, black) along mouse chromosome 17 (**H)**. Bottom image is a two Mb zoom centered on the “escaping” gene *Prdm9*. Coordinate of SPO11-oligo were obtained from Lange et al., 2016. Transcriptional changes were calculated from scRNA-seq data (ratio Le/Diff Sg1).

Though *Prdm9* ^-/-^ and *Spo11* ^-/-^ mice are defectives for DSBs localization and formation respectively, in both mutants, γH2AX is still present, albeit at low level in *Spo11* ^-/-^ spermatocytes. We thus decided to analyze the transcriptional activity in *Atm* null mutants. We used *Atm* ^-/-^;*Spo11* ^+/-^ mice as these progress up to the diplotene stage allowing the study of transcriptional arrest (EMLT) and transcriptional reactivation. In these mutants, γH2AX signaling of DSBs occurring at leptotene stage is absent (45). At leptotene and zygotene stages pPOL2 intensity was equivalent in WT and *Atm* ^-/-^;*Spo11* ^+/-^ spermatocytes (Figure 4C and D). Reactivation of transcription at pachytene stage also occurs timely (Figure 4E and F). This result indicates that DSB signaling does not influence EMLT.

In meiocytes, DSBs are rapidly repaired by homologous recombination (HR) and when global transcription is hyperactivated, few DSBs remain on autosomes from pachynemas. Meiosis specific with OB domains (MEIOB) is essential for completing meiotic recombination (22). To assess the dependence of EMLT on HR, we analyzed *Meiob* null mutants, which exhibit persistent DSBs. *Meiob*^-/-^ spermatocytes showed a clear induction of EMLT, with minimal pPOL2 staining, similarly to WT leptonema and zygonema (Figure 4C and D). In the absence of *Meiob,* prophase I arrested at early pachytene-like stage. The few mutant spermatocytes that reached this stage showed a mild elevation of pPOL2 signal, indicating that in the absence of DSB repair, transcription could be reactivated. Interestingly, pPOL2 reappeared as small patches in *Meiob*^-/-^ spermatocytes and these localized both in synapsed and unsynapsed chromosomal domains (Supplementary Figure S11B). This confirms the observation from *Hormad1* mutant revealing that synapsis is not required to terminate EMLT and to resume transcription.

Since DSBs do not appear to affect EMLT, we conjectured the reverse: that transcription might shape meiotic DSBs. Consequently, we thus tested the hypothesis that an association between recombination sites and transcriptional silencing may exist. To evaluate this hypothesis, the distance of the “silenced” genes and 1000 random intergenic regions to the nearest recombination site was computed based on DSB hotspot data defined by anti-DMC1 ssDNA sequencing (35). The distance distribution of the genes or regions to the nearest hotspots was plotted in a boxplot according to their chromosomal location (Figure 4G). The “silenced” genes were closer to the recombination sites than the random intergenic regions both on autosomes and sex chromosomes. These results suggest a possible link between distance to recombination sites and EMLT with DSBs occurring preferentially in transcriptionally silent domains. Nevertheless, given that both DSBs and silencing events occur throughout the whole genome, it was challenging to further test this relationship. Indeed, a detailed examination of a high-resolution DSB map provided by Spo11-oligonucleotides (12), revealed that distribution of DSBs and that of silenced genes largely overlapped. We exemplified this using chromosome 17 that contains several “escaping” genes including the *Prdm9* (Figure 4H). Albeit *Prdm9* is indeed located between two DSB hotspots, the presence of actively transcribed genes in leptotene is rare and is not systematic between distant hotspots.

Altogether these results indicate that EMLT is independent of the induction, signaling and repair of DSBs though it may be related to the localization of programmed DSBs.

### Stra8 regulates EMLT induction

Given that EMLT occurs around meiotic entry, we analysed pPOL2 in germ cells mutant for the main regulators of meiotic initiation. *Stra8* is a key gene involved in mitotic to meiotic transition (46, 47). It is initially expressed in A spermatogonia following their differentiation and it is strongly expressed during pre-leptotene stage. STRA8 regulates the transcription of large sets of genes. Analysis of *Stra8* null mutant mice revealed the absence of the primary induction of EMLT. In fact, quantification of pPOL2 in testis sections of Stra8^-/-^ mice showed a high level of pPOL2 in intermediate and B spermatogonia and in pre-leptotene spermatocytes when compared to corresponding stages in WT (Figure 5 and Supplementary Figure S14 and S15). MEIOC is another key factor that regulates meiosis initiation through modulation of post-transcriptional events. Contrarily to *Stra8*, genetic depletion of *Meioc*, did not modify EMLT (Figure 5 and Supplementary Figure S15), indicating that simply impairing meiotic entry was insufficient to delay the transcriptional arrest. Thus, establishment of meiotic program through STRA8 is important for the correct temporal regulation of EMLT.

**Figure 5.**
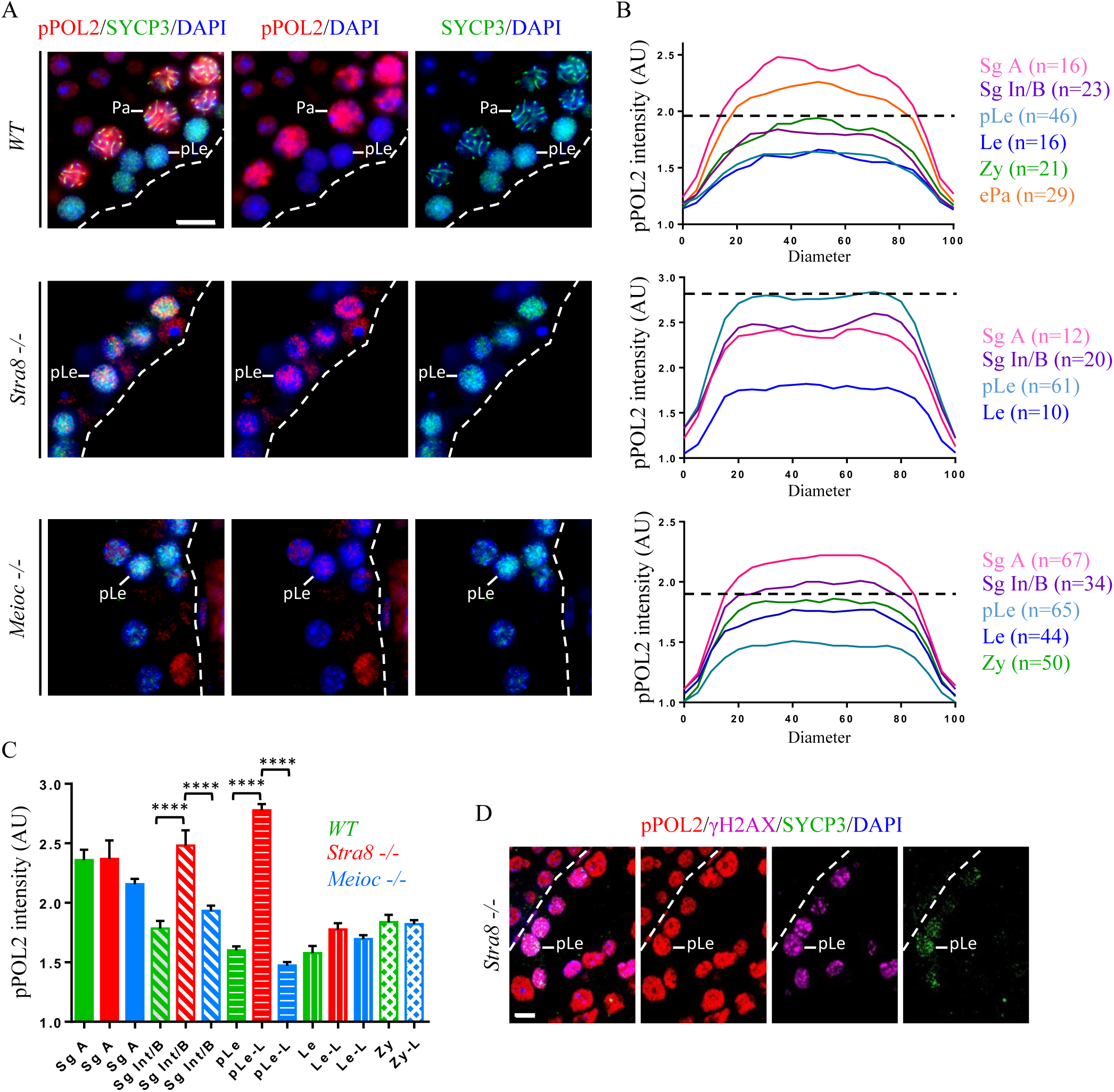
EMLT is dependent on the key regulator of mitosis-meiosis transition, STRA8. Co-immunofluorescence staining of pPOL2 (red) with SYCP3 (green) on testis sections from WT and Stra8 -/- and Meioc -/- mice (**A**). Preleptotene (pLe) cells are indicated and DNA is stained with DAPI (blue, A and D). Scale bar= 10 µm. Line plot quantification of pPOL2 intensity in WT (upper), Stra8 -/- (middle) and Meioc -/- (lower) (**B**). The horizontal axis represents the diameter of the cells. (AU=Arbitrary Units, normalization was based on background level). Legend is as in Figure 1B. The number of analysed cells is indicated in brackets for each cell type. Mean pPOL2 intensity within nuclei (**C**). Columns represent mean ± sem. In Stra8 -/-, Sg Int/B and pLe cells are more stained for pPOL2 than in WT and Meioc -/-. AU=Arbitrary Units. **** P<0.0001 (Ordinary one-way ANOVA, multiple comparisons). Co-labelling of pPOL2 (red), γH2AX (magenta) and SYCP3 (green) in Stra8 -/- testes sections (**D**). pLe cells in Stra8 -/- testes, stained for γH2AX and SYCP3, are not subjected to EMLT and show pPOL2 staining. Scale bar= 5µm.

### RNAs are massively stabilized during EMLT

The combined duration of leptotene and zygotene stage is expected to be approximately three days. How do cells survive and progress through these stages with hardly any transcription is intriguing. Comparing the “spliced” and “unspliced” expression profiles, which reflect the presence of transcripts and neo-transcription, respectively, we observed that both decreased notably at meiotic entry (Figure 6A). However, the decrease of “spliced” RNA was less pronounced indicating that some transcripts persisted. As an example, the *Rplp0* housekeeping gene displayed a twenty-fold decrease in transcription while its mRNA merely decreased two-fold during EMLT (i.e. in leptotene, unspliced mRNA equal to 4.3% versus spliced mRNA equal to 39% of the value in undifferentiated spermatogonia). We thus considered the ratio of “spliced” to “unspliced” expression levels to obtain a proxy of RNA stability. An increase of RNA stability was observed during EMLT for both “silenced” and “escaping” genes in scRNA-seq data (Figure 6A). Similar results were observed using the EU-RNA-seq dataset (Supplementary Figure S16). We hypothesized that this stabilization may be involved in launching the meiotic program and thus focused on four sets of highly expressed early meiotic genes (Figure 6B, Supplementary Table S2). These 91 genes were grouped based on their neo-transcription profile (Set 1 = early transcribed genes; Set 2 = genes transcribed during Le/Zy; Set 3 = late transcribed genes; Set 4 = genes silenced during Le/Zy). RNA stability during EMLT was also observed for the four sets of genes. These observations were also confirmed in the bulk approach with sorted cells (Supplementary Figure S17). Four well-known meiotic genes (*Dmc1 Prdm9, Meioc* and *Hormad2*), representing each set, were used to exemplify this (Figure 6C). Although, these displayed distinct neo-transcription profiles (unspliced RNA), they were robustly and simultaneously expressed in early meiosis (spliced RNA), with their stability sharply increasing in leptotene and zygotene cells. These results suggest that the global stabilisation of mRNA during EMLT and meiotic entry may facilitate the synchronised rise of meiotic gene expression.

**Figure 6.**
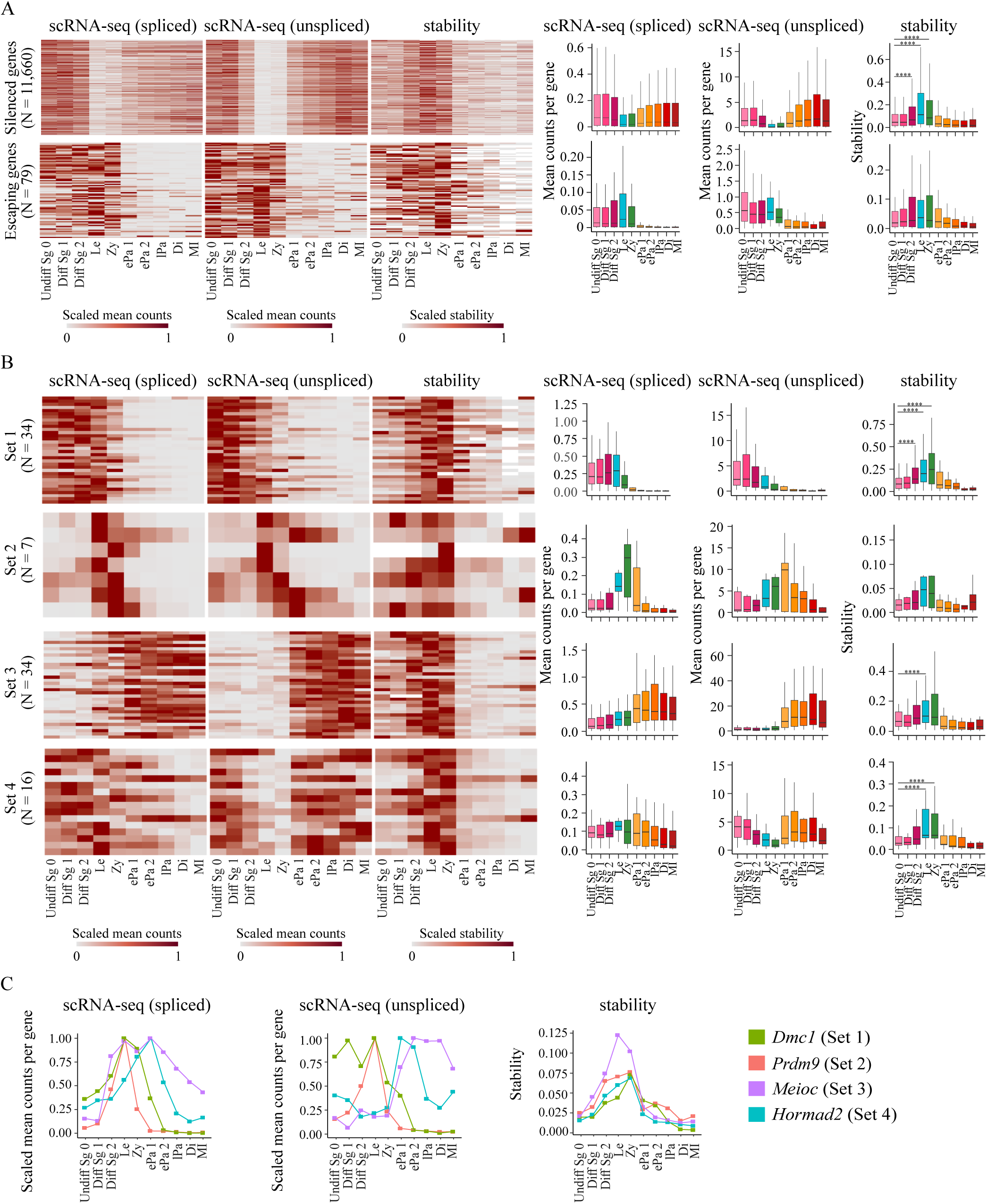
RNA stabilization occurs during EMLT. Heatmaps and boxplots of the spliced (left) and unspliced (middle) expression profiles and the stability (right) of “Silenced” (top) and “Escaping” (bottom) genes (**A**), and of four sets of early meiotic genes (**B**). Set 1 = early transcribed genes; Set 2 = genes transcribed during Le/Zy; Set 3 = late transcribed genes; Set 4 = genes silenced during Le/Zy. The “spliced” and “unspliced” expression profiles reflect transcription and neo-transcription, respectively, while the “stability” is the ratio of “spliced” to “unspliced” expression levels. In the heatmaps, each row represents a gene, and each column represents a specific spermatogenic cell population. For heatmaps, “spliced” and “unspliced” expression values of each gene were scaled between [0;1] using a 0-max scale (0=0; maximum value = 1), stability of each gene was scaled between [0;1] using a conventional scaling (minimum value = 0; maximum value = 1). Boxplots display the average counts per gene for the “spliced” and “unspliced” dataset, and stability per gene. Undiff Sg 0= Undifferentiated spermatogonia; Diff Sg = differentiated spermatogonia; Le = leptotene; Zy = zygotene; ePa = early pachytene spermatocytes; lPa = late pachytene; Di = diplotene spermatocytes; MI = Metaphase I spermatids. Gene expression and stability profiles of four representative genes from undifferentiated spermatogonia to Metaphase I spermatocytes (**C**). Plots represent the scaled average counts per gene from spliced (left) and unspliced (middle) data, and the resulting stability (right). Each of the selected genes (*Dmc1, Prdm9, Meioc* and *Hormad2*) belong to one of the four sets of meiotic genes presented in panel B.

## DISCUSSION

During the reproductive cycle, several transcriptional silencing events occur in germ cells and in the zygote (48–50). In this study, we characterized EMLT, a genome-wide silencing phenomenon during early meiotic prophase I. EMLT was dependent of the meiotic gatekeeper STRA8 but did not rely on other key events of meiotic prophase I for its establishment.

Examples of genome-wide transcriptional arrest are uncommon and notably associated with gametogenesis. Such occurrences have been well-documented post-fertilization during the early embryonic development and in primordial germ cells. The tight packaging of chromatin in the spermatozoa also induces a transcriptional silencing, though the cause is evident in this context (51). Here, we finely described a fourth major silencing event at the time of meiotic entry. The reason why germ cells are that frequently subjected to transcriptional arrests remains unclear. One may consider that among all cells, germ cells must transition between highly different states, justifying an extreme transcriptional dynamic. Indeed, germ cells need to undergo various specifications: first PGC are committed to the germ lineage; then they shift from a mitotic to a meiotic program; and later, post-meiotic male germ cells undergo a massive cellular reorganisation. Finally, after the fusion of gamete genomes, the zygote must regain totipotency. Interestingly, all these changes are all accompanied by major chromatin remodelling events, such as methylation erasure (52), meiotic recombination (53), histone to protamine transition (54) or diploidization. Of note, methylation erasure occurs in PGCs, during the first embryonic cleavages and at the time of meiotic entry (55). A simple view may accept that a quiet chromatin is a conductive environment for such events, and that an active transcriptional machinery could potentially interfere with these processes.

Recently, low transcription during leptonema and zygonema has also been reported in a study proposing it may reflect a promoter proximal paused state of POL2 (56). Another recent study described a POL2 pausing in spermatogonia (57). It is therefore conceivable that the absence of POL2 detection on chromatin in our work merely reflects a paused state.

EMLT occurring around the mitotic-meiotic switch is intriguing. Here we reported, through the analysis of several meiotic mutants, that this transcriptional arrest occurred similarly when the specific meiotic chromosomal architecture or DSB program was impaired. Cohesins shape the chromosomal architecture and have been proposed as key regulators of transcription (58–60). The involvement of specific cohesins during meiosis would be a tentative hypothesis to explain a change in transcriptional activity. However, neither *Stag3* nor *Rad21L* depletion impacted EMLT, despite the structure of meiotic chromosomes being deeply altered in such mutants. On the other hand, transcription has been proven to move cohesins (61, 62). One may thus speculate that a reverse relationship exists between EMLT and cohesins. During early meiosis, the lack of transcription may provide some stability to cohesins to initiate the axial-loop structure and the formation of the axial element formation. Another significant event occurring during early meiosis is the formation and repair of hundreds of programmed DSBs. In somatic cells, DNA damage repair has been reported to interfere with transcription in many ways, likely because both phenomena require an accessible chromatin (63). Initially, a rather straight hypothesis would have been that DSB patterning, formation, signaling or repair might trigger a transcriptional arrest. The analysis *of Prdm9, Spo11, Atm/Atr* and *Meiob* mutants contradicts such hypothesis. A reverse relationship, i.e. that active transcription might interfere with the proper generation or repair of programmed DSB, may then be considered. In this line, it is worth quoting that DSB sites are marked by H3K4me3, this histone modification is also the one retrieved in the promoters of transcriptionally active genes. A speculative assumption could be that sharing such a mark is risky and may confuse the recruitment of the DSB forming machinery. Therefore, a state of near-extinct transcription might favour the proper localisation of DSB in hotspots outside gene promoters. This hypothesis gains support from our observation that recombination occurred preferentially in transcriptionally silent regions and is in agreements with a publication indicating that DSB occur in untranscribed genomic regions (56). In the same line, it was also recently proposed that paused POL2 sites in spermatogonia correlate with those of DSBs induced by SPO11 in spermatocytes (57).

To test such hypotheses, it would be necessary to manipulate EMLT and this implies understanding its mechanisms or trigger. Here we report that EMLT is significantly delayed in absence of *Stra8*. STRA8 controls the expression of the near complete meiotic gene network. Specifically, it has recently been shown to amplify a pre-existing meiotic program (10). This aligns with the concept that STRA8 might regulate a meiotic factor that causes EMLT. In the absence of *Stra8*, such a factor would be expressed at low levels, thus delaying EMLT. Unfortunately, identifying this factor within the thousands of genes that compose the STRA8-dependent meiotic program remains a challenge. The mechanism causing EMLT also remains evasive. We report here that this mechanism differs from MSCI that is regulated by DSB signaling, asynapsis and the SETDB1 histone methylase. However, the absence of recruitment of POL2 and pPOL2 is a common feature shared by MSCI and EMLT. This reinforces the view that MSCI might be a gonosome-specific prolongation of a wider phenomenon, EMLT. In this line, it could be conceived that the histone mark repressing transcription during MSCI would also be shared with EMLT. H3K9me3 is the mark deposited by SETDB1 involved in instructing MSCI (13), though its exact role has recently been debated (16). Interestingly, H3K9me3 is also present on silent chromosomes during early prophase I, making it a likely candidate to induce EMLT. In this scenario, EMLT would rely on a different methylase which may explain that few genes on sex chromosomes are differentially regulated during EMLT and MSCI.

Gene expression can be considered as the outcome of transcription and mRNA decay rates. Our work also revealed that EMLT is accompanied by a wide event of RNA stabilisation. This stabilisation event likely buffered the decrease in transcription at meiotic onset. This aligns with a report indicating that in human cell lines, global transcriptional shutdown leads to stabilization of nearly all of transcripts through inhibition of the mRNA degradation machinery (64). Whether a similar relationship between low transcription and RNA stability exist in germ cells is a point that warrants future studies. Strikingly, when assessing the transcription of genes specifically expressed at the onset of meiosis, we observed that many (84/91) were predominantly transcribed at other stages. For most of them (sets 1, 3 and 4), their transcription decreased at the time of meiotic entry. Thus, the increased expression of these meiotic genes is mainly due changes in RNA stability. Meiotic licensing depends upon *Dazl* that is expressed early during fetal life, yet male germ cells do not initiate meiosis before post-natal life (65). This allows speculation that part of the meiotic program might be set up early, and that the decision to actually initiate meiosis requires the stabilisation of RNA to allow meiotic genes transcribed at low levels to reach a sufficient level. The second effect of this massive RNA stabilisation we observed is likely to synchronise various ‘meiotic subprograms’ to provide a timely expression of all genes needed in early meiotic prophase I. Such a hypothesis positions mRNA stabilisation as a key player orchestrating meiotic entry. In *Schizosaccharomyces pombe*, the proper timing of meiosis is controlled by the sequestration of the RNA-binding protein Mmi1, that triggers the degradation of some meiotic RNA during mitotic growth (66). In *Caenorhabditis elegans*, meiotic entry is partly controlled by a poly(A)-polymerase, GLD-2, that increases mRNA stability (67, 68). Thus, a change of RNA stability might be an ancestral mechanism controlling the mitotic-meiotic switch that has yet poorly been investigated in mammals.

In conclusion, our study provides an in-depth description of the genome-wide silencing event reported to occur during leptotene and zygotene stages. We quantified this event and proved it initiates prior to meiotic entry and is regulated by the meiotic gatekeeper, STRA8. This silencing and the accompanying RNA stabilisation are key to ensure a robust switch from the mitotic to the meiotic program. Future research will have to identify the specific mechanisms responsible for EMLT and through their manipulation it is expected that the reasons for such a major genome silencing will be unveiled.

## DATA AVAILABILITY

The data underlying this article will be shared upon request to the corresponding authors.

## SUPPLEMENTARY DATA

Supplementary Data are available online.

## ACKNOWLEDGEMENTS

The authors sincerely thank Dr J.M. Turner for helpful discussion, advices and providing the samples from *Setb1* and *Atr/Atm mutant mice*. We are indebted to Dr F. Baudat and Dr B. de Massy and Dr C. Grey for the generous provision of testicular samples from *Spo11*, *Prdm9*, and mice. Flow cytometry and cell sorting were carried out at the iRCM Flow Cytometry Shared Resource. We also thank the team within the animal housing facility at the iRCM, particularly V. Neuville and the Centre National de Recherche en Génomique Humaine for sequencing, D. Moison and S. Tourpin for her technical assistance and A. Gouret for her secretarial assistance.

## REFERENCES

1. Griswold, M.D. (2016) Spermatogenesis: The Commitment to Meiosis. Physiol Rev, 96, 1–17.

2. de Rooij, D.G. (2001) Proliferation and differentiation of spermatogonial stem cells. Reproduction, 121, 347–354.

3. Feng, C.-W., Bowles, J. and Koopman, P. (2014) Control of mammalian germ cell entry into meiosis. Mol Cell Endocrinol, 382, 488–497.

4. Hermo, L., Pelletier, R.-M., Cyr, D.G. and Smith, C.E. (2010) Surfing the wave, cycle, life history, and genes/proteins expressed by testicular germ cells. Part 2: changes in spermatid organelles associated with development of spermatozoa. Microsc Res Tech, 73, 279–319.

5. Chalmel, F. and Rolland, A.D. (2015) Linking transcriptomics and proteomics in spermatogenesis. Reproduction, 150, R149–157.

6. Margolin, G., Khil, P.P., Kim, J., Bellani, M.A. and Camerini-Otero, R.D. (2014) Integrated transcriptome analysis of mouse spermatogenesis. BMC Genomics, 15, 39.

7. Murat, F., Mbengue, N., Winge, S.B., Trefzer, T., Leushkin, E., Sepp, M., Cardoso-Moreira, M., Schmidt, J., Schneider, C., Mößinger, K., et al. (2023) The molecular evolution of spermatogenesis across mammals. Nature, 613, 308–316.

8. Naro, C., Jolly, A., Di Persio, S., Bielli, P., Setterblad, N., Alberdi, A.J., Vicini, E., Geremia, R., De la Grange, P. and Sette, C. (2017) An Orchestrated Intron Retention Program in Meiosis Controls Timely Usage of Transcripts during Germ Cell Differentiation. Dev Cell, 41, 82–93.e4.

9. Turner, J.M.A. (2015) Meiotic Silencing in Mammals. Annu Rev Genet, 49, 395–412.

10. Kojima, M.L., de Rooij, D.G. and Page, D.C. (2019) Amplification of a broad transcriptional program by a common factor triggers the meiotic cell cycle in mice. Elife, 8, e43738.

11. Cooper, T.J., Garcia, V. and Neale, M.J. (2016) Meiotic DSB patterning: A multifaceted process. Cell Cycle, 15, 13–21.

12. Lange, J., Yamada, S., Tischfield, S.E., Pan, J., Kim, S., Zhu, X., Socci, N.D., Jasin, M. and Keeney, S. (2016) The Landscape of Mouse Meiotic Double-Strand Break Formation, Processing, and Repair. Cell, 167, 695–708.e16.

13. Hirota, T., Blakeley, P., Sangrithi, M.N., Mahadevaiah, S.K., Encheva, V., Snijders, A.P., ElInati, E., Ojarikre, O.A., de Rooij, D.G., Niakan, K.K., et al. (2018) SETDB1 Links the Meiotic DNA Damage Response to Sex Chromosome Silencing in Mice. Dev Cell, 47, 645–659.e6.

14. ElInati, E., Russell, H.R., Ojarikre, O.A., Sangrithi, M., Hirota, T., de Rooij, D.G., McKinnon, P.J. and Turner, J.M.A. (2017) DNA damage response protein TOPBP1 regulates X chromosome silencing in the mammalian germ line. Proc Natl Acad Sci U S A, 114, 12536–12541.

15. Royo, H., Prosser, H., Ruzankina, Y., Mahadevaiah, S.K., Cloutier, J.M., Baumann, M., Fukuda, T., Höög, C., Tóth, A., de Rooij, D.G., et al. (2013) ATR acts stage specifically to regulate multiple aspects of mammalian meiotic silencing. Genes Dev, 27, 1484–1494.

16. Abe, H., Yeh, Y.-H., Munakata, Y., Ishiguro, K.-I., Andreassen, P.R. and Namekawa, S.H. (2022) Active DNA damage response signaling initiates and maintains meiotic sex chromosome inactivation. Nat Commun, 13, 7212.

17. Abby, E., Tourpin, S., Ribeiro, J., Daniel, K., Messiaen, S., Moison, D., Guerquin, J., Gaillard, J.-C., Armengaud, J., Langa, F., et al. (2016) Implementation of meiosis prophase I programme requires a conserved retinoid-independent stabilizer of meiotic transcripts. Nat Commun, 7, 10324.

18. Monesi, V. (1964) RIBONUCLEIC ACID SYNTHESIS DURING MITOSIS AND MEIOSIS IN THE MOUSE TESTIS. J Cell Biol, 22, 521–532.

19. Page, J., de la Fuente, R., Manterola, M., Parra, M.T., Viera, A., Berríos, S., Fernández-Donoso, R. and Rufas, J.S. (2012) Inactivation or non-reactivation: what accounts better for the silence of sex chromosomes during mammalian male meiosis? Chromosoma, 121, 307–326.

20. Hogarth, C.A., Evanoff, R., Mitchell, D., Kent, T., Small, C., Amory, J.K. and Griswold, M.D. (2013) Turning a spermatogenic wave into a tsunami: synchronizing murine spermatogenesis using WIN 18, 446. Biol Reprod, 88, 40.

21. Bastos, H., Lassalle, B., Chicheportiche, A., Riou, L., Testart, J., Allemand, I. and Fouchet, P. (2005) Flow cytometric characterization of viable meiotic and postmeiotic cells by Hoechst 33342 in mouse spermatogenesis. Cytometry Part A, 65A, 40–49.

22. Souquet, B., Abby, E., Hervé, R., Finsterbusch, F., Tourpin, S., Le Bouffant, R., Duquenne, C., Messiaen, S., Martini, E., Bernardino-Sgherri, J., et al. (2013) MEIOB targets single-strand DNA and is necessary for meiotic recombination. PLoS Genet, 9, e1003784.

23. Widger, A., Mahadevaiah, S.K., Lange, J., ElInati, E., Zohren, J., Hirota, T., Pacheco, S., Maldonado-Linares, A., Stanzione, M., Ojarikre, O., et al. (2018) ATR is a multifunctional regulator of male mouse meiosis. Nat Commun, 9, 2621.

24. Diagouraga, B., Clément, J.A.J., Duret, L., Kadlec, J., de Massy, B. and Baudat, F. (2018) PRDM9 Methyltransferase Activity Is Essential for Meiotic DNA Double-Strand Break Formation at Its Binding Sites. Mol Cell, 69, 853–865.e6.

25. Daniel, K., Lange, J., Hached, K., Fu, J., Anastassiadis, K., Roig, I., Cooke, H.J., Stewart, A.F., Wassmann, K., Jasin, M., et al. (2011) Meiotic homologue alignment and its quality surveillance are controlled by mouse HORMAD1. Nat Cell Biol, 13, 599–610.

26. Herrán, Y., Gutiérrez-Caballero, C., Sánchez-Martín, M., Hernández, T., Viera, A., Barbero, J.L., de Álava, E., de Rooij, D.G., Suja, J.Á., Llano, E., et al. (2011) The cohesin subunit RAD21L functions in meiotic synapsis and exhibits sexual dimorphism in fertility. EMBO J, 30, 3091–3105.

27. Llano, E., Gomez-H, L., García-Tuñón, I., Sánchez-Martín, M., Caburet, S., Barbero, J.L., Schimenti, J.C., Veitia, R.A. and Pendas, A.M. (2014) STAG3 is a strong candidate gene for male infertility. Hum Mol Genet, 23, 3421–3431.

28. Dereli, I., Stanzione, M., Olmeda, F., Papanikos, F., Baumann, M., Demir, S., Carofiglio, F., Lange, J., de Massy, B., Baarends, W.M., et al. (2021) Four-pronged negative feedback of DSB machinery in meiotic DNA-break control in mice. Nucleic Acids Res, 49, 2609–2628.

29. Ernst, C., Eling, N., Martinez-Jimenez, C.P., Marioni, J.C. and Odom, D.T. (2019) Staged developmental mapping and X chromosome transcriptional dynamics during mouse spermatogenesis. Nat Commun, 10, 1251.

30. McCarthy, D.J., Campbell, K.R., Lun, A.T.L. and Wills, Q.F. (2017) Scater: pre-processing, quality control, normalization and visualization of single-cell RNA-seq data in R. Bioinformatics, 33, 1179– 1186.

31. McGinnis, C.S., Murrow, L.M. and Gartner, Z.J. (2019) DoubletFinder: Doublet Detection in Single-Cell RNA Sequencing Data Using Artificial Nearest Neighbors. Cell Syst, 8, 329–337.e4.

32. Barh, D. and Azevedo, V.A.D.C. (2019) Single-cell Omics: Technological Advances and Applications Academic Press Inc, LondonLJ; San Diego, CA.

33. Stuart, T., Butler, A., Hoffman, P., Hafemeister, C., Papalexi, E., Mauck, W.M., Hao, Y., Stoeckius, M., Smibert, P. and Satija, R. (2019) Comprehensive Integration of Single-Cell Data. Cell, 177, 1888–1902.e21.

34. La Manno, G., Soldatov, R., Zeisel, A., Braun, E., Hochgerner, H., Petukhov, V., Lidschreiber, K., Kastriti, M.E., Lönnerberg, P., Furlan, A., et al. (2018) RNA velocity of single cells. Nature, 560, 494– 498.

35. Smagulova, F., Brick, K., Pu, Y., Camerini-Otero, R.D. and Petukhova, G.V. (2016) The evolutionary turnover of recombination hot spots contributes to speciation in mice. Genes Dev, 30, 266–280.

36. Quinlan, A.R. and Hall, I.M. (2010) BEDTools: a flexible suite of utilities for comparing genomic features. Bioinformatics, 26, 841–842.

37. Chen, Y., Lyu, R., Rong, B., Zheng, Y., Lin, Z., Dai, R., Zhang, X., Xie, N., Wang, S., Tang, F., et al. (2020) Refined spatial temporal epigenomic profiling reveals intrinsic connection between PRDM9-mediated H3K4me3 and the fate of double-stranded breaks. Cell Res, 30, 256–268.

38. Teletin, M., Vernet, N., Ghyselinck, N.B. and Mark, M. (2017) Roles of Retinoic Acid in Germ Cell Differentiation. Curr Top Dev Biol, 125, 191–225.

39. Llano, E., Herrán, Y., García-Tuñón, I., Gutiérrez-Caballero, C., de Álava, E., Barbero, J.L., Schimenti, J., de Rooij, D.G., Sánchez-Martín, M. and Pendás, A.M. (2012) Meiotic cohesin complexes are essential for the formation of the axial element in mice. J Cell Biol, 197, 877–885.

40. Hamer, G., Roepers-Gajadien, H.L., van Duyn-Goedhart, A., Gademan, I.S., Kal, H.B., van Buul, P.P.W. and de Rooij, D.G. (2003) DNA double-strand breaks and gamma-H2AX signaling in the testis. Biol Reprod, 68, 628–634.

41. Baudat, F., Buard, J., Grey, C., Fledel-Alon, A., Ober, C., Przeworski, M., Coop, G. and de Massy, B. (2010) PRDM9 is a major determinant of meiotic recombination hotspots in humans and mice. Science, 327, 836–840.

42. Parvanov, E.D., Petkov, P.M. and Paigen, K. (2010) Prdm9 controls activation of mammalian recombination hotspots. Science, 327, 835.

43. Baudat, F., Manova, K., Yuen, J.P., Jasin, M. and Keeney, S. (2000) Chromosome synapsis defects and sexually dimorphic meiotic progression in mice lacking Spo11. Mol Cell, 6, 989–998.

44. Romanienko, P.J. and Camerini-Otero, R.D. (2000) The mouse Spo11 gene is required for meiotic chromosome synapsis. Mol Cell, 6, 975–987.

45. Barchi, M., Mahadevaiah, S., Di Giacomo, M., Baudat, F., de Rooij, D.G., Burgoyne, P.S., Jasin, M. and Keeney, S. (2005) Surveillance of different recombination defects in mouse spermatocytes yields distinct responses despite elimination at an identical developmental stage. Mol Cell Biol, 25, 7203– 7215.

46. Anderson, E.L., Baltus, A.E., Roepers-Gajadien, H.L., Hassold, T.J., de Rooij, D.G., van Pelt, A.M.M. and Page, D.C. (2008) Stra8 and its inducer, retinoic acid, regulate meiotic initiation in both spermatogenesis and oogenesis in mice. Proc Natl Acad Sci U S A, 105, 14976–14980.

47. Koubova, J., Menke, D.B., Zhou, Q., Capel, B., Griswold, M.D. and Page, D.C. (2006) Retinoic acid regulates sex-specific timing of meiotic initiation in mice. Proc Natl Acad Sci U S A, 103, 2474–2479.

48. Schulz, K.N. and Harrison, M.M. (2019) Mechanisms regulating zygotic genome activation. Nat Rev Genet, 20, 221–234.

49. Seki, Y., Yamaji, M., Yabuta, Y., Sano, M., Shigeta, M., Matsui, Y., Saga, Y., Tachibana, M., Shinkai, Y. and Saitou, M. (2007) Cellular dynamics associated with the genome-wide epigenetic reprogramming in migrating primordial germ cells in mice. Development, 134, 2627–2638.

50. Bonnerot, C., Vernet, M., Grimber, G., Briand, P. and Nicolas, J.F. (1991) Transcriptional selectivity in early mouse embryos: a qualitative study. Nucleic Acids Res, 19, 7251–7257.

51. Sassone-Corsi, P. (2002) Unique chromatin remodeling and transcriptional regulation in spermatogenesis. Science, 296, 2176–2178.

52. Sanz, L.A., Kota, S.K. and Feil, R. (2010) Genome-wide DNA demethylation in mammals. Genome Biol, 11, 110.

53. Baudat, F., Imai, Y. and de Massy, B. (2013) Meiotic recombination in mammals: localization and regulation. Nat Rev Genet, 14, 794–806.

54. Hoghoughi, N., Barral, S., Vargas, A., Rousseaux, S. and Khochbin, S. (2018) Histone variants: essential actors in male genome programming. J Biochem, 163, 97–103.

55. Huang, Y., Li, L., An, G., Yang, X., Cui, M., Song, X., Lin, J., Zhang, X., Yao, Z., Wan, C., et al. (2023) Single-cell multi-omics sequencing of human spermatogenesis reveals a DNA demethylation event associated with male meiotic recombination. Nat Cell Biol, 25, 1520–1534.

56. Alexander, A.K., Rice, E.J., Lujic, J., Simon, L.E., Tanis, S., Barshad, G., Zhu, L., Lama, J., Cohen, P.E. and Danko, C.G. (2023) A-MYB and BRDT-dependent RNA Polymerase II pause release orchestrates transcriptional regulation in mammalian meiosis. Nat Commun, 14, 1753.

57. Kaye, E.G., Basavaraju, K., Nelson, G.M., Zomer, H.D., Roy, D., Joseph, I.I., Rajabi-Toustani, R., Qiao, H., Adelman, K. and Reddi, P.P. (2024) RNA polymerase II pausing is essential during spermatogenesis for appropriate gene expression and completion of meiosis. Nat Commun, 15, 848.

58. Braccioli, L. and de Wit, E. (2019) CTCF: a Swiss-army knife for genome organization and transcription regulation. Essays Biochem, 63, 157–165.

59. Yuen, K.C. and Gerton, J.L. (2018) Taking cohesin and condensin in context. PLoS Genet, 14, e1007118.

60. Wang, J., Bando, M., Shirahige, K. and Nakato, R. (2022) Large-scale multi-omics analysis suggests specific roles for intragenic cohesin in transcriptional regulation. Nat Commun, 13, 3218.

61. Lengronne, A., Katou, Y., Mori, S., Yokobayashi, S., Kelly, G.P., Itoh, T., Watanabe, Y., Shirahige, K. and Uhlmann, F. (2004) Cohesin relocation from sites of chromosomal loading to places of convergent transcription. Nature, 430, 573–578.

62. Borrie, M.S., Campor, J.S., Joshi, H. and Gartenberg, M.R. (2017) Binding, sliding, and function of cohesin during transcriptional activation. Proc Natl Acad Sci U S A, 114, E1062–E1071.

63. Khobta, A. and Epe, B. (2012) Interactions between DNA damage, repair, and transcription. Mutat Res, 736, 5–14.

64. Slobodin, B., Bahat, A., Sehrawat, U., Becker-Herman, S., Zuckerman, B., Weiss, A.N., Han, R., Elkon, R., Agami, R., Ulitsky, I., et al. (2020) Transcription Dynamics Regulate Poly(A) Tails and Expression of the RNA Degradation Machinery to Balance mRNA Levels. Mol Cell, 78, 434–444.e5.

65. Lin, Y., Gill, M.E., Koubova, J. and Page, D.C. (2008) Germ cell-intrinsic and -extrinsic factors govern meiotic initiation in mouse embryos. Science, 322, 1685–1687.

66. Harigaya, Y., Tanaka, H., Yamanaka, S., Tanaka, K., Watanabe, Y., Tsutsumi, C., Chikashige, Y., Hiraoka, Y., Yamashita, A. and Yamamoto, M. (2006) Selective elimination of messenger RNA prevents an incidence of untimely meiosis. Nature, 442, 45–50.

67. Kadyk, L.C. and Kimble, J. (1998) Genetic regulation of entry into meiosis in Caenorhabditis elegans. Development, 125, 1803–1813.

68. Nousch, M., Yeroslaviz, A., Habermann, B. and Eckmann, C.R. (2014) The cytoplasmic poly(A) polymerases GLD-2 and GLD-4 promote general gene expression via distinct mechanisms. Nucleic Acids Res, 42, 11622–11633.

