## Supplemental Tables, Figures and Legends for "Genome-wide transcriptional silencing and mRNA stabilization allow the coordinated expression of the meiotic program in mice"

**Legend to Supplementary Tables:**

**Table S1. List of primary and secondary antibodies used in this article for immunofluorescence (IF).**
Poly, Polyclonal antibody; Mono, Monoclonal antibody.

**Table S2. List of genes analysed in Figure 6 B.**
These genes are highly expressed in early meiotic prophase I.

**Legend to Supplementary Figures:**

**Supp. Fig. S1. At Pachytene stage sex body is subjected to Meiotic Sex Chromosome Inactivation.**
Co-immunofluorescence staining for POL2 (red), pPOL2 (green) and SYCP3 (blue) on chromosome spreads in leptotene (Le), early- and mid-pachytene (ePa, mPa) spermatocytes (**A**). Orange dashed squares represent the sex body.
Plot profile quantification of pPOL2 (left) and POL2 (right). Each line represents one representative cell (**B**). pLe = preleptotene; Le = leptotene; mPa = mid-pachytene. The horizontal axis indicates the cell diameter. For each point along the line, the intensity of pPOL2 or POL2 is quantified with ImageJ software. MSCI is indicated with an orange dashed line.

**Supp. Fig. S2.**
For each cell, intensity of pPOL2 (left) and POL2 (right) is quantified in testis sections with ImageJ software thanks to Plot profile analysis (**A**). Each line represents a cell. Cells are ordered top-down from the less differentiated to the more differentiated stage of spermatogenesis. Sg A= Type A spermatogonia, n=19; Sg Int/B= intermediate and type B spermatogonia, n=26; pLe= preleptotene, n=27; Le=leptotene, n=27; Zy=zygotene, n=22; ePa=early pachytene, n=27; mPa=mid pachytene, n=50; lPa/Di=late pachytene/diplotene, n=32; rSd=round spermatids, n=50; eSd=elongated spermatids, n=4. Each column represents a point along the diameter of the cell, from 0 to 10 (with 0 and 10 representing the cytoplasm of the cell and 5 representing the centre of the cell). ePa spermatocytes (orange rectangle) are low for pPOL2 and high for POL2.
Quantification of pPOL2 (upper graph) and POL2 (below) intensity in nuclei (**B**). Histograms represent means ± sem for each cell type. Different letters indicate significant differences (ordinary one-way ANOVA, multiple comparisons).

**Supp. Fig. S3. Correlation between POL2 and pPOL2.**
POL2 and pPOL2 intensities were measured as described in Supp 2. Left panel represents all spermatogenic stages. Right panels focus on the changes occurring from type A spermatogonia to leptotene spermatocytes (top), from early pachytene to late pachytene/diplotene (middle) and from round to elongated spermatids (bottom). These illustrate the setup of EMLT, the hyperactivation of POL2 and the transcriptional repression occurring during spermiogenesis respectively.

**Supp. Fig. S4. Workflows for bioinformatics analyses.**
Spliced and unspliced analysis of the mouse testis single-cell atlas (**A**). Mouse scRNA-seq fastq files were downloaded from EBI (E-MTAB-

6946). These included ten samples from mice aged 5 to 42 postnatal days (PND5 to 42). Raw sequencing reads were processed and mapped to the mouse (mm10) genome with *Cell Ranger* (v4) with default parameters. This output was used i) to generate unspliced count data with the *Velocyto* tool, and ii) for *Seurat* analysis after quality controls with *Scater*, *Scran* and *DoubletFinder*. Batch effect correction on the samples was carried out with CCA prior to data normalization. Dimensionality reduction was performed by with PCA followed by UMAP. Cells were clustered based on the PCA data and the resulting cell clusters were annotated based on known marker genes. Cell clusters corresponding to germ cell populations were subsetted and submitted to the same pipeline as described above. Finally, a ploidy correction was performed on the unspliced count matrix prior to its integration into a Seurat object.
Identification of "silenced" and "escaping" genes (**B**). The ploidy-corrected unspliced count matrix was first normalized for intron size. Genes showing a null variance were next discarded. A two-step process was then used to identify potential "silenced" and "escaping" genes. Genes were considered "silenced" if their expression values were: *i)* higher than an expression threshold (median value of the overall count matrix =0.07) in at least one of the relevant stages (Diff Spg 1, Diff Spg 2, ePa 1, ePa 2, lPa and/or Di) and, ii) significantly correlated (p-value < 0.05) to the expression profiles of archetypal genes (*Ndufa11*, *Park7*) showing a silenced expression pattern in Le and Zy. Genes were considered "escaping" if their expression values were: i) higher than 0.07 in Le and Zy; and, ii) significantly correlated (p-value < 0.05) to archetypal genes (*Pet2*, *Ccnb3*, *Prdm9*) showing an escaping expression pattern in Le or Zy.
Processing of the bulk RNA-seq and EU-RNA-seq datasets (**C**). RNA-seq and EU-RNA-seq raw reads were mapped onto the mouse genome (mm10) using STAR. The *StringTie* tool was used for transcript assembly while *Ballgown* was used for quantification. The resulting FPKM expression matrix from the EU-RNA-seq dataset was corrected by the expression of 5S ribosomal genes which are considered to be stable and independent of POL2 (ie. transcribed by RNA POL3).

**Supp. Fig. S5. Characterization of mouse testicular cells.**
Mouse testis scRNA-seq data from Ernst et al., 2019 were downloaded (EBI : E-MTAB-6946) and analysed as described in Supplementary figure S4.
Bar plots of the number of cells and of the median number of detected genes per cell for each sample (**A**). Each sample corresponds to a distinct colour from post-natal day (PND) 5 to the adult stage.
UMAP plots before (**B**) and after (**C**) batch effect correction of all cells from pooled samples coloured according to panel A. Deconvolution of the global batch effect corrected UMAP from PND5 to adult stage (**D**). The cells are coloured by sample according to the colour code in panel A.
UMAP plot, clustering and annotation of all testicular cells (**E**).
Dotplot showing the expression of gene markers that were used for cell cluster annotation (**F**). The size of a dot corresponds to the percentage of cells expressing a given gene marker while its colour represents the corresponding scaled average expression.

**Supp. Fig. S6. Characterization of germ cells using scRNA-seq data.**

UMAP plot of all germ cells from post-natal day (PND) 5 to the adult stage from the Ernst et al. (2019) dataset( **A**). Each cluster corresponds to a distinct spermatogenic stage. Undiff Sg = Undifferentiated spermatogonia; Diff Sg = Differentiated spermatogonia; Le = leptotene spermatocytes; Zy = zygotene spermatocytes; ePa = early pachytene spermatocytes; lPa = late pachytene spermatocytes; Di = diplotene spermatocytes; MI = Metaphase I spermatocytes; MII = Metaphase II spermatocytes; rSd = round spermatids, eSd = elongated spermatids.
Dotplot showing the overlap between cell clusters from the current analysis and those from Ernst et al. (2019) (**B, Left**). The size of a dot corresponds to the percentage of cells from a given cluster (this analysis) associated to the original annotation (Ernst et al., 2019). The colour represents the scaled -log10 (p-value) of the enrichment test. Dotplot showing the expression of gene markers in germ cell clusters from the current analysis (**B, Middle**). The size of a dot corresponds to the percentage of cells expressing a given gene while its colour represents the corresponding scaled average expression. A stacked bar plot showing the cell cycle phase assigned to the cells (G1, S or G2/M) for each cluster (**B, right**).
Boxplots of transcriptional activity from autosomes or sex chromosomes during spermatogenesis before (top) and after (bottom) ploidy correction split by autosomes or sex chromosomes (**C**). Boxplots represent counts per cells from each spermatogenic stage and are coloured according to panel A.

**Supp. Fig. S7. Synchronisation of spermatogenesis for EU-RNA-seq.**
Protocol for EU-RNA-seq (**A**). Spermatogenetic waves were synchronised as previously described by Hogarth et al (2013). Two days post-partum (dpp) mice were pipette-fed daily with WIN 18,446 and at 9 dpp animals were injected with retinoic acid (RA). At 24 dpp, mice were injected with Ethynyl-Uridine (EU). Three hours later mice were sacrificed, testes were dissociated, and cells were FACS-sorted based on their ploidy and red Hoechst exclusion. Spermatogonia (Sg), Leptotene (Le) and Pachytene (Pa) spermatocytes were used for total RNA sequencing while nascent RNAs were captured using Click-iT capture and sequenced (EU-RNA-Seq).
Immunofluorescence staining for pPOL2 (red) and γH2AX (green) in 24 dpp synchronised versus non-synchronised testes (**B**). Stages of the seminiferous epithelium are indicated with Roman numerals. Tubules are delineated by dotted lines. DNA is stained with DAPI (blue).

**Supp. Fig. S8. Conservation of EMLT in female mice and other mammals.**
Immunofluorescence staining of pPOL2 (red) on testicular sections from different mammalian species (Goat, Rabbit and Dog) (**A**). Spermatocytes are identified thanks to γH2AX (green) staining and DNA is stained with DAPI (blue). At the beginning of prophase I (diffuse γH2AX, white arrows) pPOL2 is very low. At the end of prophase I (γH2AX at sex body, white stars) pPOL2 staining is higher. Dotted white curves delimit seminiferous tubules. Scale bars= 10μm.
Characterisation of pPOL2 expression profile during prophase I in female mice by co-immunolabelling of pPOL2 (green) and SYCP3 (red) on chromosome spreads (**B**). DNA is stained with DAPI (blue). At each stage two cells are presented, one with low pPOL2 staining and one with

abundant staining. Le=leptotene; Zy=zygotene; Pa=pachytene; Di=diplotene.
Quantification of pPOL2 intensity on mouse oocyte chromosome spreads (**C**). pPOL2 intensity is normalised to mitotic cells (100%, black dashed line). Each violet dot represents one cell. Oocytes were retrieved from different embryos (n=5); black bars represent means ± sem. pPOL2 staining intensity is heterogeneous at each stage of prophase I. lDi=late diplotene. Different letters indicate significant differences (Ordinary one-way ANOVA, multiple comparisons).

**Supp. Fig. S9. Analysis of silenced and escaping gene expression in bulk and single cells.**
Heatmap of the expression of the 11,660 "silenced" genes distributed according to their chromosomal location (autosomes and sex chromosomes) from the unspliced scRNA-seq (average count matrix) and from the EU-RNA-seq dataset (FPKM matrix) (**A**). These datasets reflect neo-transcription, and are used to study transcriptional activity during spermatogenesis. In the heatmaps, each row represents a gene, and each column represents a specific spermatogenic cell population (scRNA-seq) or a sample (EU-RNA-seq). For heatmaps, expression values (average counts for unspliced scRNA-seq and FPKM for EU-RNA-seq) of each gene were scaled between [0;1] using a 0-max scale (0=0; maximum value = 1). Diff Sg = differentiated spermatogonia; Le = leptotene spermatocytes; Zy = zygotene spermatocytes; ePa = early pachytene spermatocytes; lPa = late pachytene spermatocytes; Di = diplotene spermatocytes; Sg = spermatogonia; Pa = pachytene spermatocytes.
Heatmap of the expression of the 79 "escaping" genes distributed according to their chromosomal location (autosomes and sex chromosomes), from the unspliced scRNA-seq (average count matrix) and the EU-RNA-seq dataset (FPKM matrix), respectively (**B**). Figure legends are the same as in panel A.

**Supp. Fig. S10. Quantification of histone modifications on silenced and escaping genes.**
Quantification of seven histone modifications on the 11,660 "silenced" and 79 "escaping" genes according to ChIP-seq data from Chen et al. (2020; GSE132446) (**A**).
Enrichment profiles of "silenced" genes (blue), "escaping" genes (red), and "random regions" (grey) (**B**). Values represent means ± sem Figure legends of heatmaps and enrichment profiles are the same as in Figure 2B.

**Supp. Fig. S11. Reappearance of pPOL2 on chromosome axes.**
Co-immunolabelling for pPOL2 (green) and SYCP3 (red) on chromosome spreads from spermatocytes. DNA is stained with DAPI (blue).
In WT pachytene spermatocytes pPOL2 reappears first on chromosome axes as patches (**A**). 90° rotation of green colour (pPOL2) confirms that patch localisation on axes is not random.
In *Hormad1* and *Meiob* mutants pPol2 reappears on synapsed as well as on unsynapsed axes at pachytene stage. White dashed squares represent enlarged images (right).

**Supp. Fig. S12. Dynamic of pPOL2 during spermatogenesis is correlated with γH2AX.**
Immunofluorescence staining of pPOL2 (red), SYCP3 (green) and γH2AX (magenta) on testes sections. DNA is stained with DAPI (blue). Type A

spermatogonia (Sg A), negative for γH2AX, are highly positive for pPOL2. Intermediate/type B spermatogonia (Sg Int/B), preleptotene (pLe) and zygotene (Zy) cells are stained for γH2AX and show a low level of pPOL2 staining. γH2AX is absent (excepted in the sex body) in late prophase I spermatocytes (mPa, lPa, Di), which are highly stained for pPOL2. Roman numerals indicate the stage of the seminiferous epithelium cycle. Dotted white curbs delimit seminiferous tubules. Scale bar= 5μm

**Supp. Fig. S13. In *Spo11 -/-* mutants EMLT is more pronounced.**
Co-immunolabelling for pPOL2 (green) and for SYCP3 (red) in *Spo11* -/- testis sections (**A**). DNA is stained with DAPI (blue). pPOL2 intensity is quantified in WT and mutant mice (**B**). SgA= Type A spermatogonia ; pLe= preleptotene; Le= leptotene; Zy=zygotene; ePa= early pachytene; -L = -like. *P<0,05 **P<0,01 **** P<0.0001 (unpaired t-test).

**Supp. Fig. S14. EMLT induction in *Stra8 -/-* mice is delayed.**
Co-immunolabelling for pPOL2 (red) and for SYCP3 (green) in *Stra8-/-* testis sections (**A**). Leptotene (Le) cells in *Stra8-/-* testis are subjected to EMLT and show weak pPOL2 staining. DNA is stained with DAPI (blue). Dotted white curbs delimit seminiferous tubules. Scale bar= 10 μm.
Co-immunolabelling of POL2 (red) and ZBTB16 (green) in *Stra8-/-* testis sections (**B**). Both type A spermatogonia (Sg A) and intermediate type and B spermatogonia (Sg Int/B) are robustly stained for pPOL2. Scale bar= 10 μm.

**Supp. Fig. S15. Dynamic of pPOL2 during spermatogenesis is different in *Stra8-/-* when compared to WT or *Meioc-/-*.**
Each line represents a different germ cell. Germ cells are ordered top-down from the less differentiated to the more differentiated stage. Sg A = Spermatogonia A; Sg Int/B = Intermediate and B Spermatogonia; pLe = preLeptotene; Le = Leptotene; Zy = Zygotene; ePa = early Pachytene. For each cell, pPOL2 intensity is measured with ImageJ software using Line Plot analysis. Each column represents a point of the diameter of the cell, from 0 to 10 (with 0 and 10 representing the cytoplasm of the cell and 5 representing the centre of the cell). pPOL2 intensity in Sg Int/B (violet rectangles) and pLe cells (light blue rectangles) in *Stra8 -/-* is higher than in *WT* and *Meioc -/-.*

**Supp. Fig. S16. Comparison of silenced and escaping genes stability between single-cell and bulk datasets.**
Heatmaps and boxplots of the expression and stability of the 10,576 "silenced" genes and 69 "escaping" genes distributed according to their chromosomal location (autosomes and sex chromosomes) from the scRNA-seq dataset (average count matrix) and the RNA-seq dataset (FPKM matrix). In the scRNA-seq dataset the "spliced" and "unspliced" expression profiles reflect transcription and neo-transcription, respectively, while the "stability" is the ratio of "spliced" to "unspliced" expression levels. In the RNA-seq dataset "RNA-seq" and "EU-RNA-seq" expression profiles reflect transcription and neo-transcription, respectively, while "stability" is the ratio of "RNA-seq" to "EU-RNA-seq" expression levels.
In the heatmaps, each row represents a gene, and each column represents a specific spermatogenic cell population(scRNA-seq) or a sample (RNA-

seq and EU-RNA-seq). For heatmaps, "RNA-seq", "EU-RNA-seq", "spliced" and "unspliced" expression values of each gene were scaled between [0;1] using a 0-max scale (0=0; maximum value = 1). The stability of each gene was scaled between [0;1] using a conventional scaling (minimum value = 0; maximum value = 1). Boxplots display the average counts per gene and the FPKM values for the scRNA-seq dataset and for RNA-seq dataset, respectively, and the stability per gene. One-tailed Wilcoxon tests were used to compare the stability per spermatogenic cell in the different sets of genes. Undiff Sg = undifferentiated spermatogonia; Diff Sg = differentiated spermatogonia; Le = leptotene spermatocytes; Zy = zygotene spermatocytes; ePa = early pachytene spermatocytes; lPa = late pachytene spermatocytes; Di = diplotene spermatocytes; MI = Metaphase I spermatids, Sg = spermatogonia; Pa = pachytene spermatocytes.

**Supp. Fig. S17. Stability of early meiotic genes in the bulk dataset.** Heatmaps and boxplots of the RNA-seq (left) and EU-RNA-seq (middle) expression profiles and of the stability (right) for four sets of early meiotic genes (Set 1 = early transcribed genes; Set 2 = genes expressed during Le/Zy; Set 3 = late transcribed genes; Set 4 = genes silenced during Le/Zy). The "RNA-seq" and "EU-RNA-seq" expression profiles reflect transcription and neo-transcription, respectively, while "stability" is the ratio of "RNA-seq" to "EU-RNA-seq" expression levels. In the heatmaps, each row represents a gene and each column represents a sample. For heatmaps, "RNA-seq" and "EU-RNA-seq" expression values of each gene were scaled between [0;1] using a 0-max scale (0=0; maximum value = 1), stability of each gene was scaled between [0;1] using a conventional scaling (minimum value = 0; maximum value = 1). Boxplots display the FPKM per gene for the "RNA-seq" and "EU-RNA-seq" datasets, and the stability per gene. Sg = spermatogonia; Le = leptotene spermatocytes; Pa = pachytene spermatocytes. One-tailed Wilcoxon tests were used to compare the stability per spermatogenic cell for different sets of genes: ****: $p < 0.0001$.

| Antibody | Host species | Mono/poly clonal | Application | Concentration | Compagny | Reference |
|---|---|---|---|---|---|---|
| POL2 | Mouse | Mono | IF | 1/200 | ThermoFisher | MAI-46093 |
| pPOL2 | Rabbit | Mono | IF | 1/200 | Abcam | ab193468 |
| STRA8 | Rabbit | Poly | IF | 1/500 | Abcam | ab49602 |
| SYCP3 | Rabbit | Poly | IF | 1/500 | Novus | NB300-232 |
| SYCP3 | Mouse | Mono | IF | 1/500 | Abcam | ab97672 |
| SYCP3 | Guinnea-Pig | Poly | IF | 1/500 | Our lab | |
| ZBTB16 | Mouse | Mono | IF | 1/200 | Santa Cruz | SC28319 |
| γH2AX | Rabbit | Poly | IF | 1/500 | Abcam | ab11174 |
| γH2AX | Mouse | Mono | IF | 1/500 | Millipore | 05-636 |
| H3K9me3 | Rabbit | Poly | IF | 1/200 | Millipore | 07-442 |
| Alexa488 anti-rabbit | Donkey | Poly | IF | 1/500 | Life technologies | A21206 |
| Alexa488 anti-mouse | Donkey | Poly | IF | 1/500 | Life technologies | A21202 |
| Alexa488 anti-guinneaPig | Goat | Poly | IF | 1/500 | Life technologies | A11073 |
| Alexa594 anti-rabbit | Donkey | Poly | IF | 1/500 | Life technologies | A21207 |
| Alexa594 anti-mouse | Donkey | Poly | IF | 1/500 | Life technologies | A21203 |
| 594 anti-guinneaPig | Donkey | Poly | IF | 1/500 | Sigma-Aldrich | SAB4600096 |
| Alexa555 anti-rabbit | Goat | Poly | IF | 1/500 | Life technologies | A21429 |
| Alexa555 anti-mouse | Goat | Poly | IF | 1/500 | Life technologies | A11003 |
| Alexa647 anti-rabbit | Donkey | Poly | IF | 1/500 | RD systems | NL005 |
| Alexa647anti-mouse | Donkey | Poly | IF | 1/500 | RD systems | NL008 |
| Alexa350 anti-guinnea-pig | Donkey | Poly | IF | 1/200 | Life technologies | SA5-10093 |

**Table S1.** List of primary and secondary antibodies used in this article.

Table S2: List of genes analysed in Figure 6 B; Genes are selected as highly expressed in early meiosis. Genes are categorised in four sets based on their transcriptional profile (unspliced RNA).

| *Name* | *Set* | *Meiosis Status* |
|---|---|---|
| 4933427D06Rik | Set 1 | early |
| Adarb1 | Set 1 | early |
| 1700013H16Rik | Set 1 | early |
| Caprin2 | Set 1 | early |
| 4930447C04Rik | Set 1 | early |
| M1ap | Set 1 | early |
| Ddb2 | Set 1 | early |
| **Dmc1** | Set 1 | early |
| Ecsit | Set 1 | early |
| Figla | Set 1 | early |
| Ccnb3 | Set 1 | early |
| Haus8 | Set 1 | early |
| Phka2 | Set 1 | early |
| Fmr1nb | Set 1 | early |
| Btbd18 | Set 1 | early |
| Rad21l | Set 1 | early |
| Stra8 | Set 1 | early |
| Rbpms2 | Set 1 | early |
| Rec8 | Set 1 | early |
| Rhox13 | Set 1 | early |
| Ribc1 | Set 1 | early |
| Zcwpw1 | Set 1 | early |
| Syn2 | Set 1 | early |
| Smc1b | Set 1 | early |
| Taf7l | Set 1 | early |
| Taf9b | Set 1 | early |
| Tex11 | Set 1 | early |
| Tex15 | Set 1 | early |
| Tktl1 | Set 1 | early |
| Tsc22d3 | Set 1 | early |
| Ugt8a | Set 1 | early |
| Fmr1 | Set 1 | early |
| Taf4b | Set 1 | early |
| Wbp2nl | Set 1 | early |
| Meiob | Set 2 | escaper |
| Fbxo47 | Set 2 | escaper |
| Pet2 | Set 2 | escaper |
| **Prdm9** | Set 2 | escaper |
| Rad51ap2 | Set 2 | escaper |
| Inca1 | Set 2 | escaper |
| Aspa | Set 3 | late |
| Majin | Set 3 | late |
| Abhd18 | Set 3 | late |
| 4930432K21Rik | Set 3 | late |
| Asf1b | Set 3 | late |
| Ccdc181 | Set 3 | late |
| 4930524B15Rik | Set 3 | late |
| BC051142 | Set 3 | late |
| Ccdc73 | Set 3 | late |

| *Name* | *Set* | *Meiosis Status* |
|---|---|---|
| Terb1 | Set 3 | late |
| Ccdc155 | Set 3 | late |
| Cntd1 | Set 3 | late |
| Dmrtc2 | Set 3 | late |
| Fhl4 | Set 3 | late |
| **Meioc** | Set 3 | late |
| Gpat2 | Set 3 | late |
| Gpr19 | Set 3 | late |
| Hsf2bp | Set 3 | late |
| Hormad1 | Set 3 | late |
| Il18 | Set 3 | late |
| Larp1b | Set 3 | late |
| Lypd4 | Set 3 | late |
| Spata22 | Set 3 | late |
| Stag3 | Set 3 | late |
| Spdya | Set 3 | late |
| Spo11 | Set 3 | late |
| Syce1 | Set 3 | late |
| Sycp2 | Set 3 | late |
| Sycp3 | Set 3 | late |
| Syngr4 | Set 3 | late |
| Tex101 | Set 3 | late |
| Tsga10 | Set 3 | late |
| Zfp541 | Set 3 | late |
| Ly6k | Set 3 | late |
| Sirt7 | Set 4 | silenced |
| Terb2 | Set 4 | silenced |
| Usp32 | Set 4 | silenced |
| BC049762 | Set 4 | silenced |
| Ccdc36 | Set 4 | silenced |
| Ccdc172 | Set 4 | silenced |
| Crebl2 | Set 4 | silenced |
| Dennd4a | Set 4 | silenced |
| Pramel1 | Set 4 | silenced |
| Spata5 | Set 4 | silenced |
| Cdkl2 | Set 4 | silenced |
| Spryd3 | Set 4 | silenced |
| **Hormad2** | Set 4 | silenced |
| Slc25a31 | Set 4 | silenced |
| Setdb2 | Set 4 | silenced |

Supp. Fig. S1

A

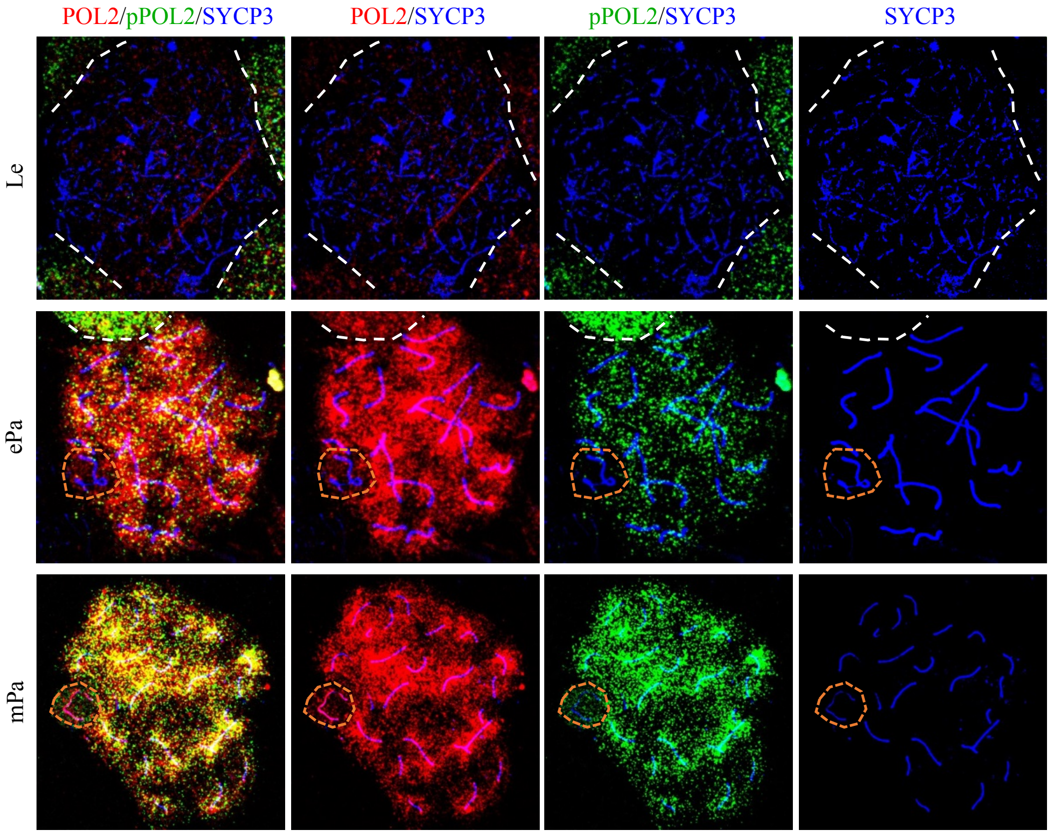

B

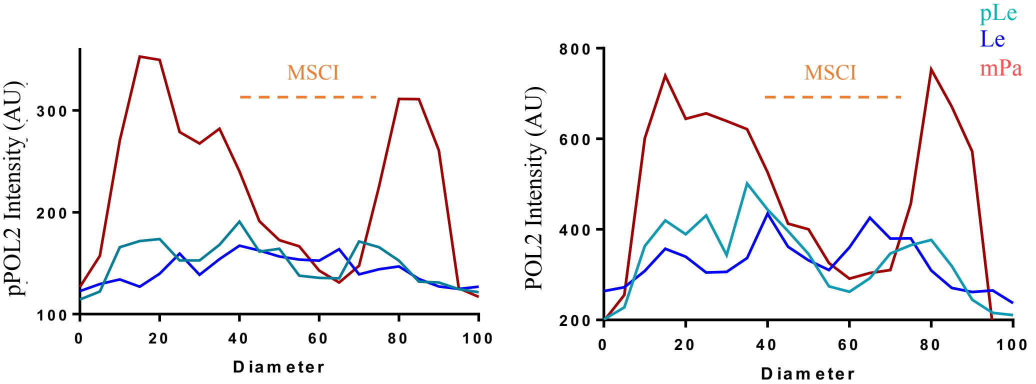

Supp. Fig. S2

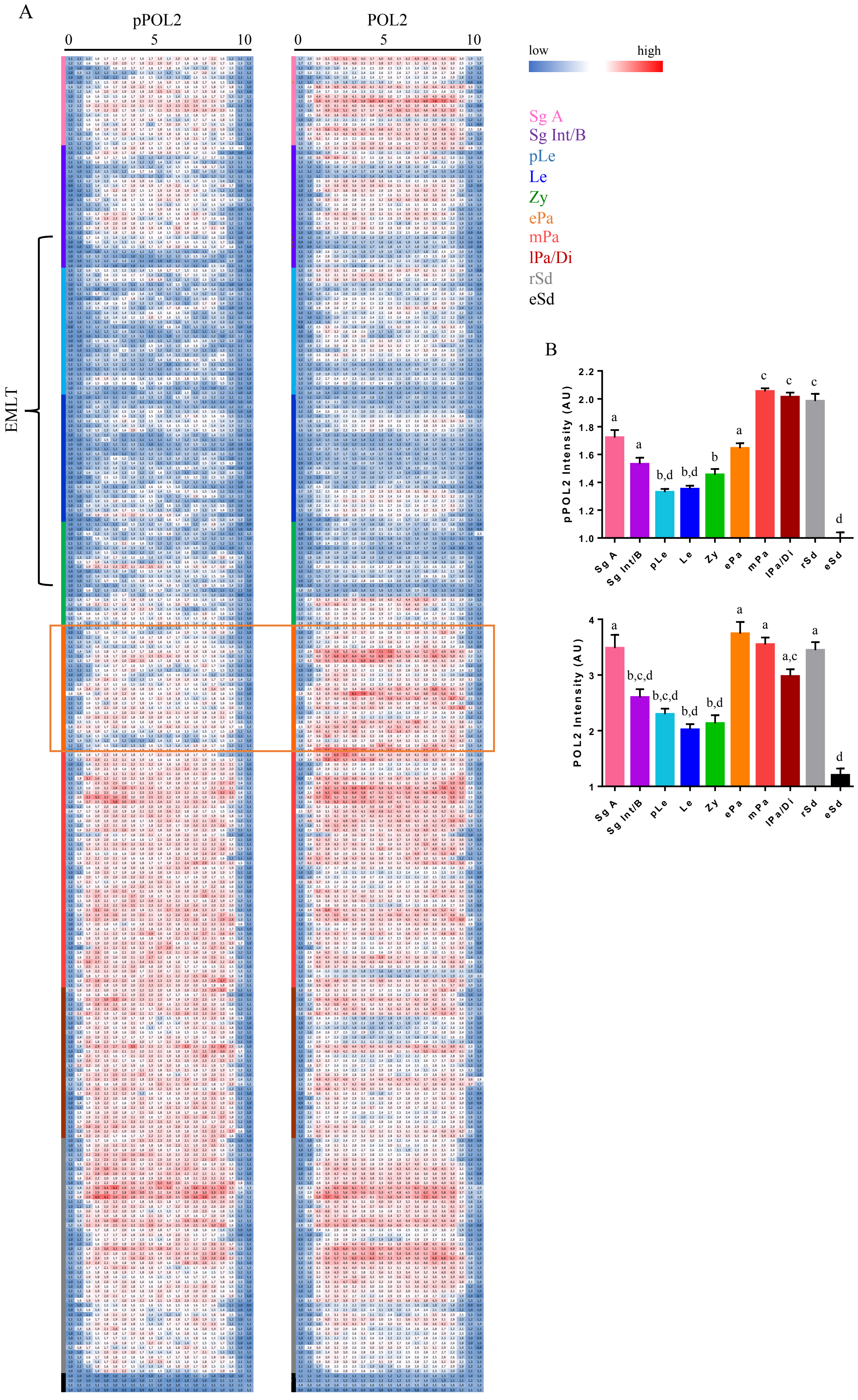
A
pPOL2
POL2
0
5
10
EMLT
low
high
Sg A
Sg Int/B
pLe
Le
Zy
ePa
mPa
lPa/Di
rSd
eSd
B
pPOL2 Intensity (AU)
1.0
1.2
1.4
1.6
1.8
2.0
2.2
a
a
b,d
b,d
b
a
c
c
c
d
POL2 Intensity (AU)
1
2
3
4
a
b,c,d
b,c,d
b,d
b,d
a
a
a,c
a
d
Sg A
Sg Int/B
pLe
Le
Zy
ePa
mPa
lPa/Di
rSd
eSd

Supp. Fig. S3

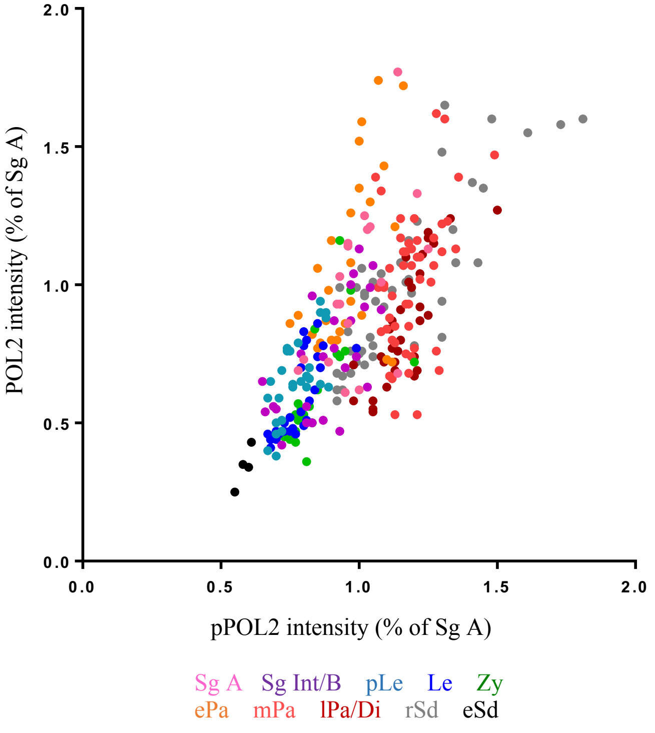
POL2 intensity (% of Sg A)
pPOL2 intensity (% of Sg A)
Sg A Sg Int/B pLe Le Zy
ePa mPa lPa/Di rSd eSd

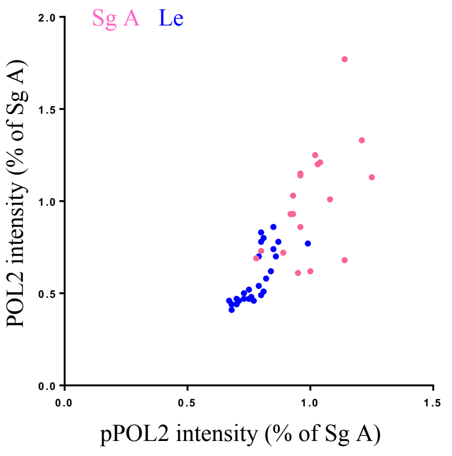
Sg A Le
POL2 intensity (% of Sg A)
pPOL2 intensity (% of Sg A)

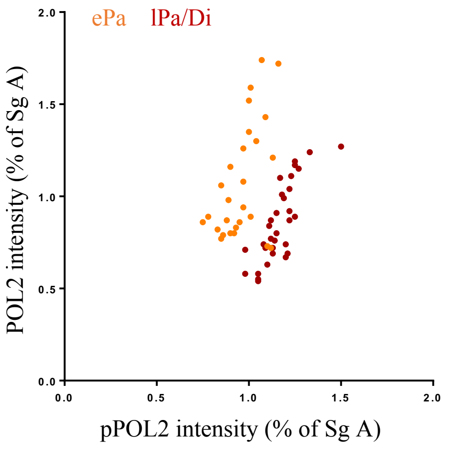
ePa lPa/Di
POL2 intensity (% of Sg A)
pPOL2 intensity (% of Sg A)

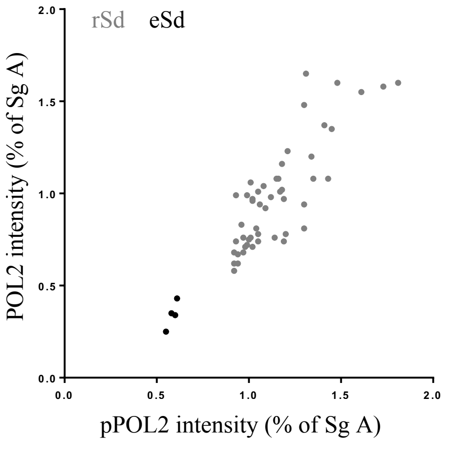
rSd eSd
POL2 intensity (% of Sg A)
pPOL2 intensity (% of Sg A)

Supp. Fig. S4

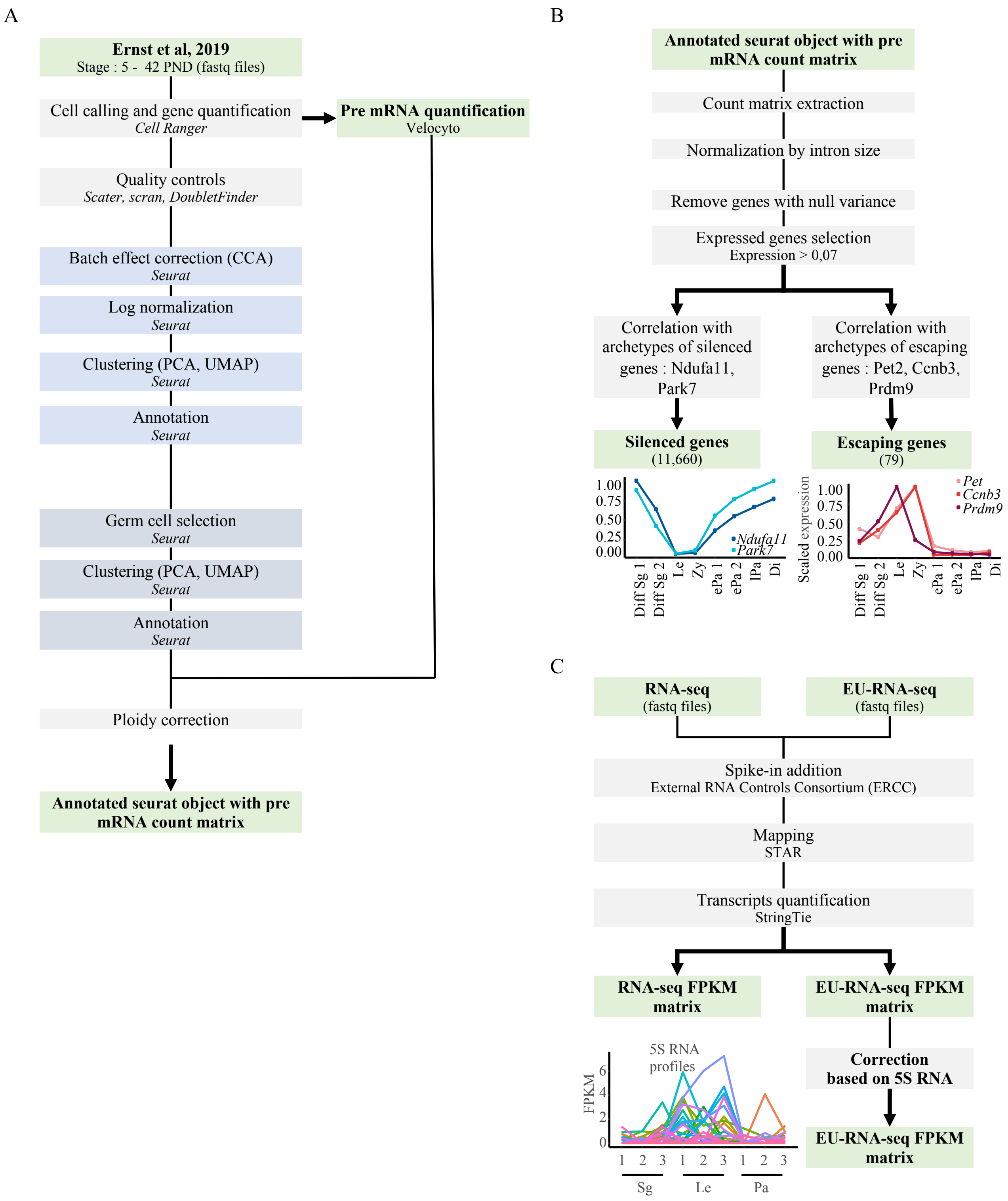
A
Ernst et al, 2019
Stage : 5 - 42 PND (fastq files)
Cell calling and gene quantification
Cell Ranger
Pre mRNA quantification
Velocyto
Quality controls
Scater, scran, DoubletFinder
Batch effect correction (CCA)
Seurat
Log normalization
Seurat
Clustering (PCA, UMAP)
Seurat
Annotation
Seurat
Germ cell selection
Seurat
Clustering (PCA, UMAP)
Seurat
Annotation
Seurat
Ploidy correction
Annotated seurat object with pre mRNA count matrix
B
Annotated seurat object with pre mRNA count matrix
Count matrix extraction
Normalization by intron size
Remove genes with null variance
Expressed genes selection
Expression > 0,07
Correlation with archetypes of silenced genes : Ndufa11, Park7
Correlation with archetypes of escaping genes : Pet2, Ccnb3, Prdm9
Silenced genes
(11,660)
Escaping genes
(79)
1.00
0.75
0.50
0.25
0.00
Ndufa11
Park7
Diff Sg 1
Diff Sg 2
Le
Zy
ePa 1
ePa 2
lPa
Di
Scaled expression
Pet
Ccnb3
Prdm9
C
RNA-seq
(fastq files)
EU-RNA-seq
(fastq files)
Spike-in addition
External RNA Controls Consortium (ERCC)
Mapping
STAR
Transcripts quantification
StringTie
RNA-seq FPKM matrix
EU-RNA-seq FPKM matrix
Correction based on 5S RNA
EU-RNA-seq FPKM matrix
5S RNA profiles
FPKM
0
2
4
6
1 2 3 1 2 3 1 2 3
Sg
Le
Pa

### Supp. Fig. S5

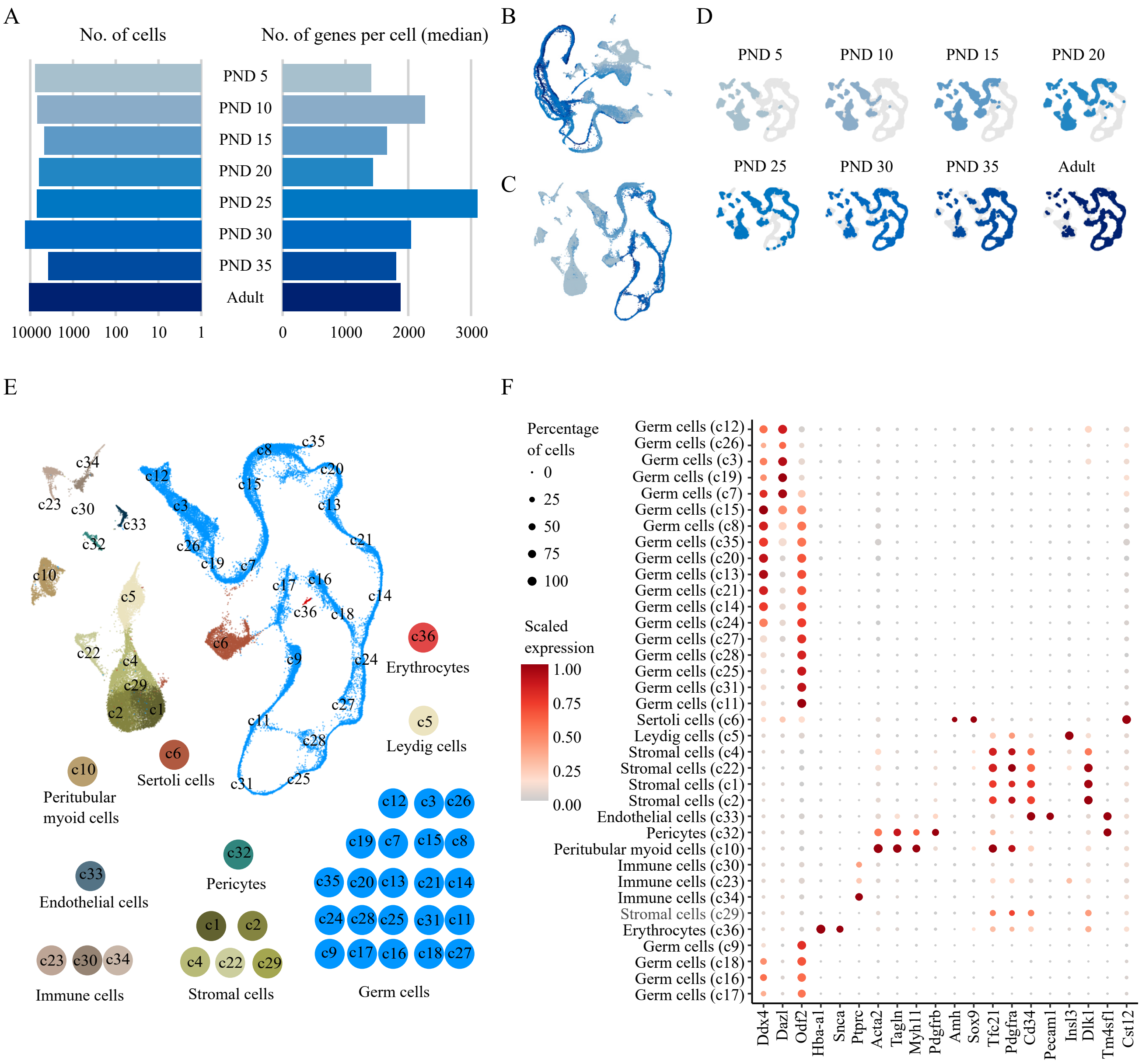

A
No. of cells
No. of genes per cell (median)
PND 5
PND 10
PND 15
PND 20
PND 25
PND 30
PND 35
Adult
10000 1000 100 10 1
0 1000 2000 3000
B
C
D
PND 5
PND 10
PND 15
PND 20
PND 25
PND 30
PND 35
Adult
E
Erythrocytes
Leydig cells
Peritubular myoid cells
Sertoli cells
Endothelial cells
Pericytes
Immune cells
Stromal cells
Germ cells
F
Percentage of cells
0
25
50
75
100
Scaled expression
1.00
0.75
0.50
0.25
0.00
Germ cells (c12)
Germ cells (c26)
Germ cells (c3)
Germ cells (c19)
Germ cells (c7)
Germ cells (c15)
Germ cells (c8)
Germ cells (c35)
Germ cells (c20)
Germ cells (c13)
Germ cells (c21)
Germ cells (c14)
Germ cells (c24)
Germ cells (c27)
Germ cells (c28)
Germ cells (c25)
Germ cells (c31)
Germ cells (c11)
Sertoli cells (c6)
Leydig cells (c5)
Stromal cells (c4)
Stromal cells (c22)
Stromal cells (c1)
Stromal cells (c2)
Endothelial cells (c33)
Pericytes (c32)
Peritubular myoid cells (c10)
Immune cells (c30)
Immune cells (c23)
Immune cells (c34)
Stromal cells (c29)
Erythrocytes (c36)
Germ cells (c9)
Germ cells (c18)
Germ cells (c16)
Germ cells (c17)
Ddx4
Dazl
Odf2
Hba-a1
Snca
Ptprc
Acta2
Tagln
Myh11
Pdgfrb
Amh
Sox9
Tfc21
Pdgfra
Cd34
Pecam1
Insl3
Dlk1
Tm4sf1
Cst12

Supp. Fig. S6

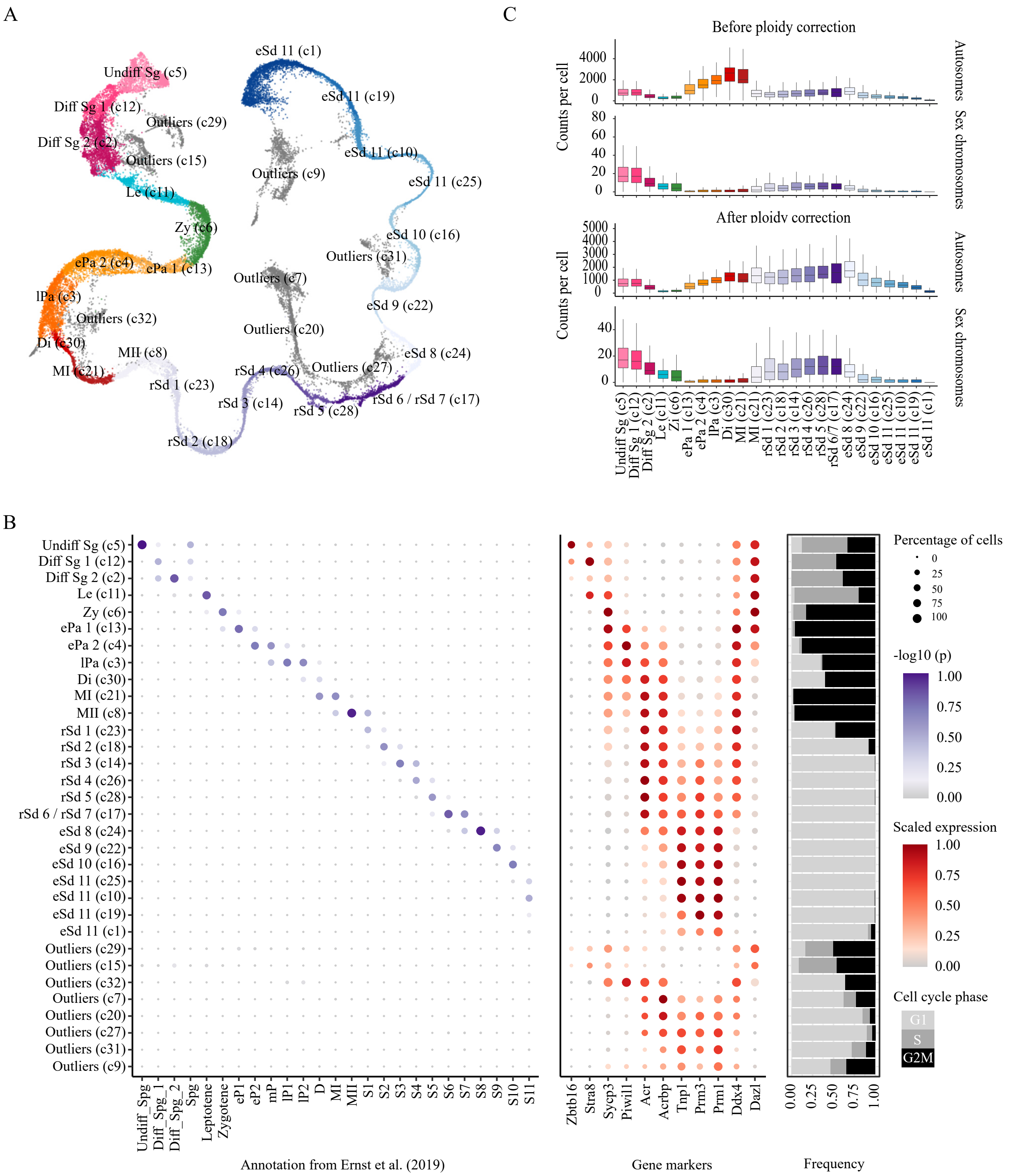
A
Undiff Sg (c5)
Diff Sg 1 (c12)
Diff Sg 2 (c2)
Outliers (c29)
Outliers (c15)
Le (c11)
Zy (c6)
ePa 2 (c4)
ePa 1 (c13)
lPa (c3)
Outliers (c32)
Di (c30)
MI (c21)
MII (c8)
rSd 1 (c23)
rSd 2 (c18)
rSd 3 (c14)
rSd 4 (c26)
rSd 5 (c28)
rSd 6 / rSd 7 (c17)
eSd 11 (c1)
eSd 11 (c19)
eSd 11 (c10)
eSd 11 (c25)
eSd 10 (c16)
eSd 9 (c22)
eSd 8 (c24)
Outliers (c9)
Outliers (c31)
Outliers (c7)
Outliers (c20)
Outliers (c27)
C
Before ploidy correction
After ploidy correction
Counts per cell
Autosomes
Sex chromosomes
B
Annotation from Ernst et al. (2019)
Undiff_Spg
Diff_Spg_1
Diff_Spg_2
Spg
Leptotene
Zygotene
eP1
eP2
mP
lP1
lP2
D
MI
MII
S1
S2
S3
S4
S5
S6
S7
S8
S9
S10
S11
Gene markers
Zbtb16
Stra8
Sycp3
Piwil1
Acr
Acrbp
Tnp1
Prm3
Prm1
Ddx4
Dazl
Frequency
Percentage of cells
-log10 (p)
Scaled expression
Cell cycle phase
G1
S
G2M

Supp. Fig. S7

A

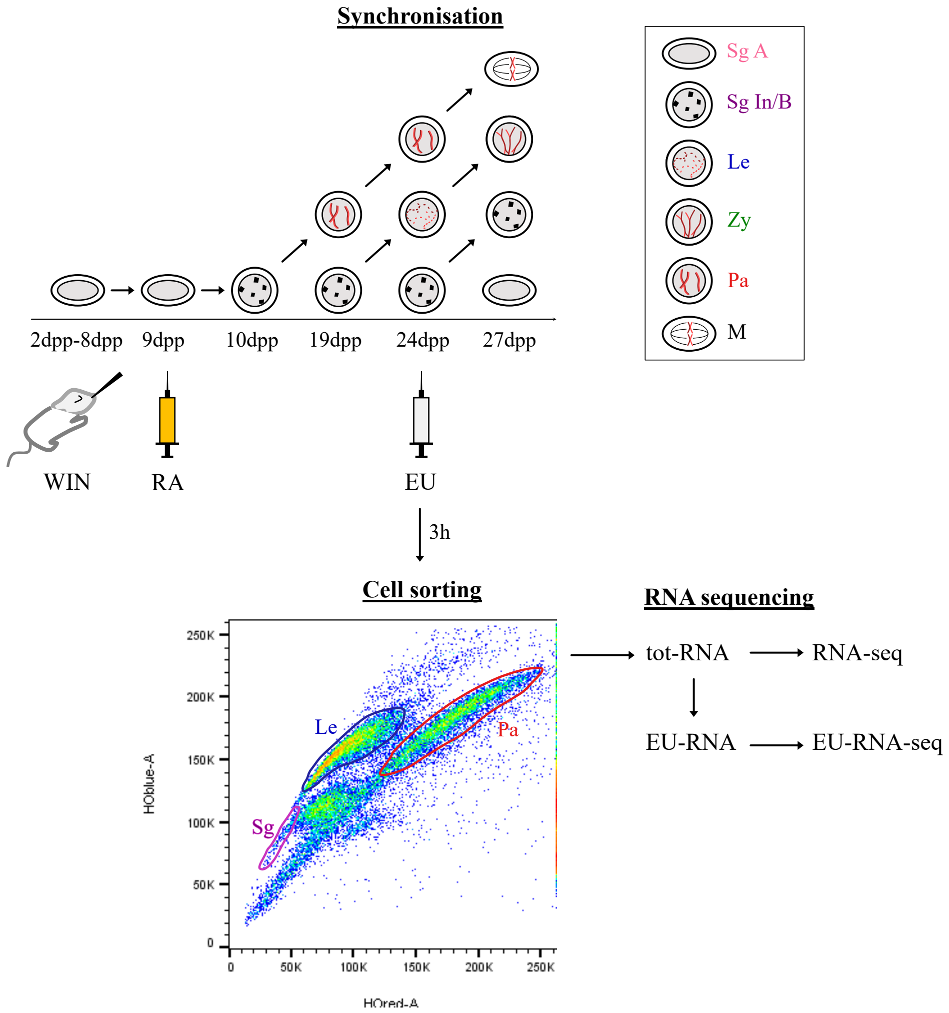

B

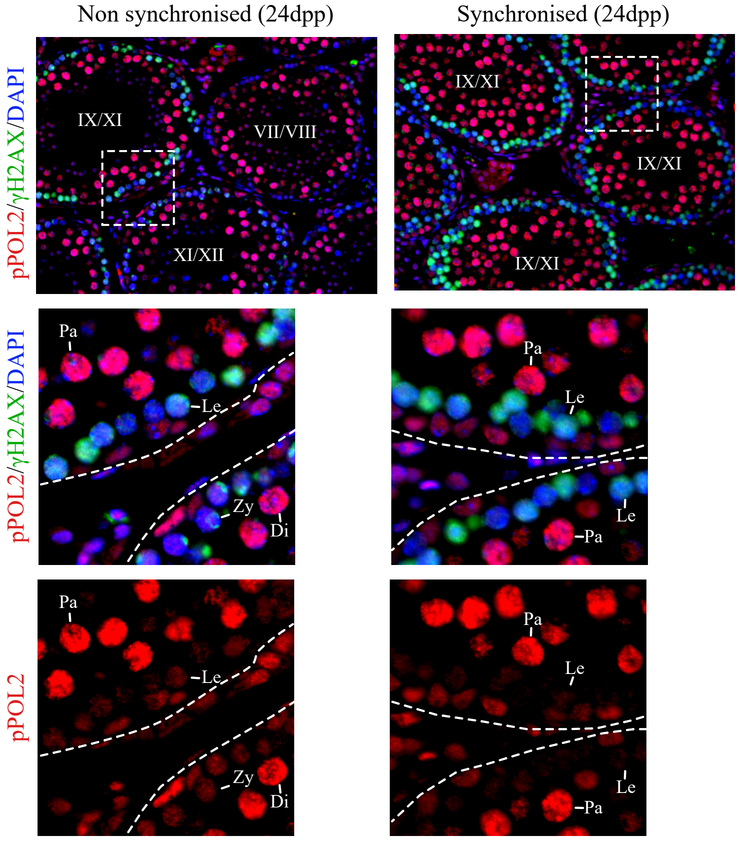

Supp. Fig. S8

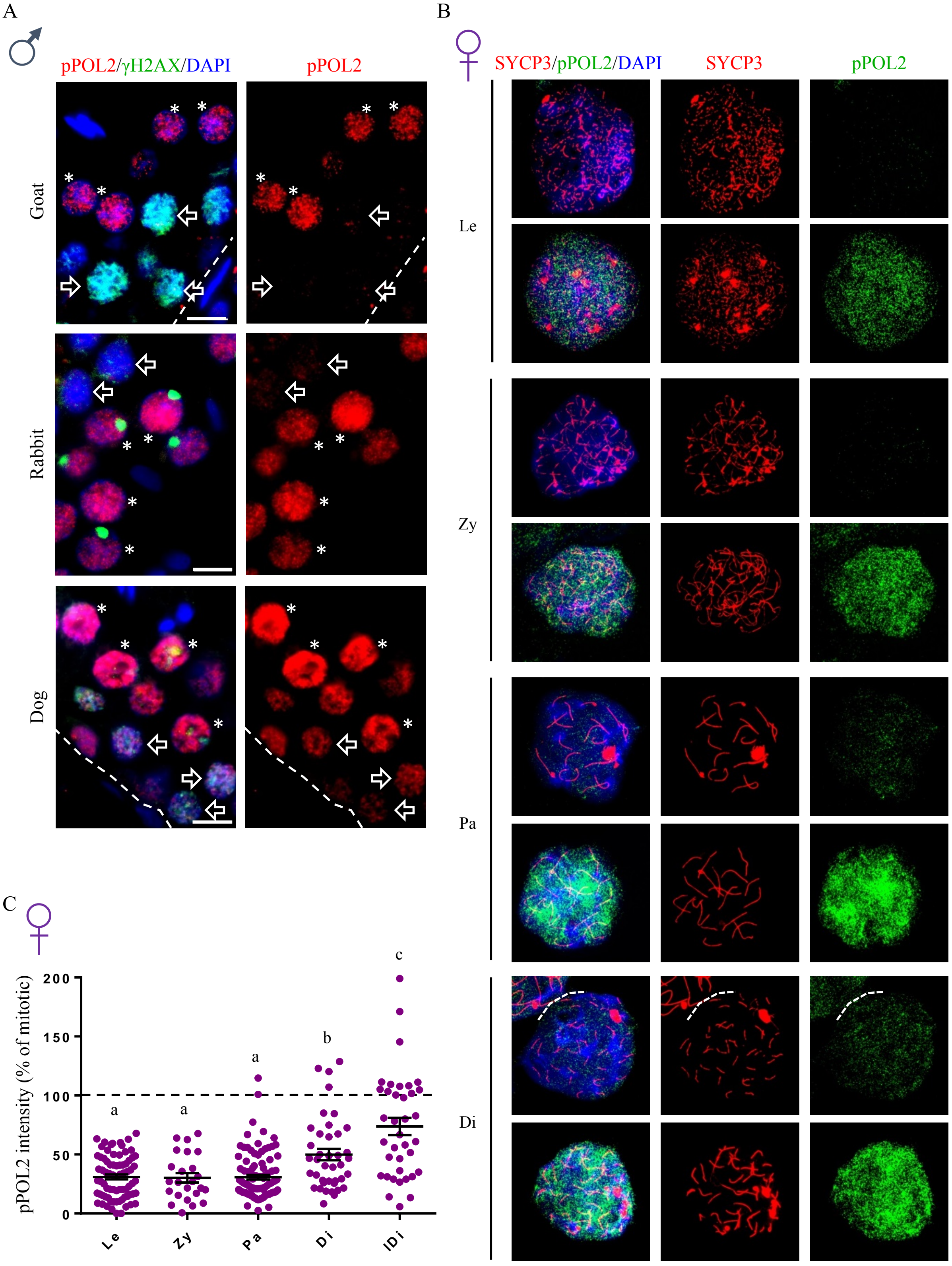

A
♂
pPOL2/γH2AX/DAPI
pPOL2
Goat
Rabbit
Dog
B
♀
SYCP3/pPOL2/DAPI
SYCP3
pPOL2
Le
Zy
Pa
Di
C
♀
pPOL2 intensity (% of mitotic)
200
150
100
50
0
a
a
a
b
c
Le
Zy
Pa
Di
lDi

Supp. Fig. S9

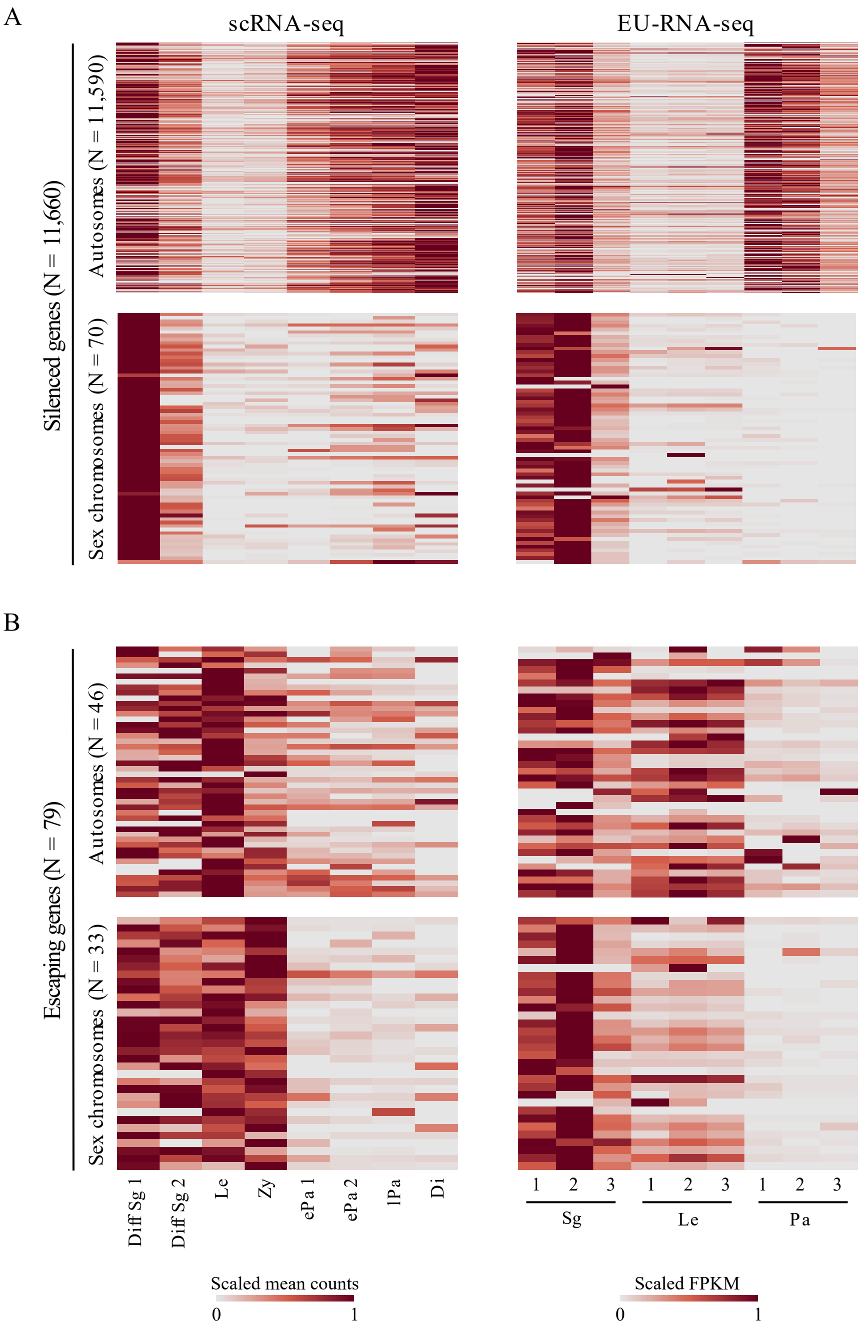
A
scRNA-seq
EU-RNA-seq
Silenced genes (N = 11,660)
Autosomes (N = 11,590)
Sex chromosomes (N = 70)
B
Escaping genes (N = 79)
Autosomes (N = 46)
Sex chromosomes (N = 33)
Diff Sg 1
Diff Sg 2
Le
Zy
ePa 1
ePa 2
lPa
Di
1 2 3 1 2 3 1 2 3
Sg
Le
Pa
Scaled mean counts
0 1
Scaled FPKM
0 1

Supp. Fig. S10

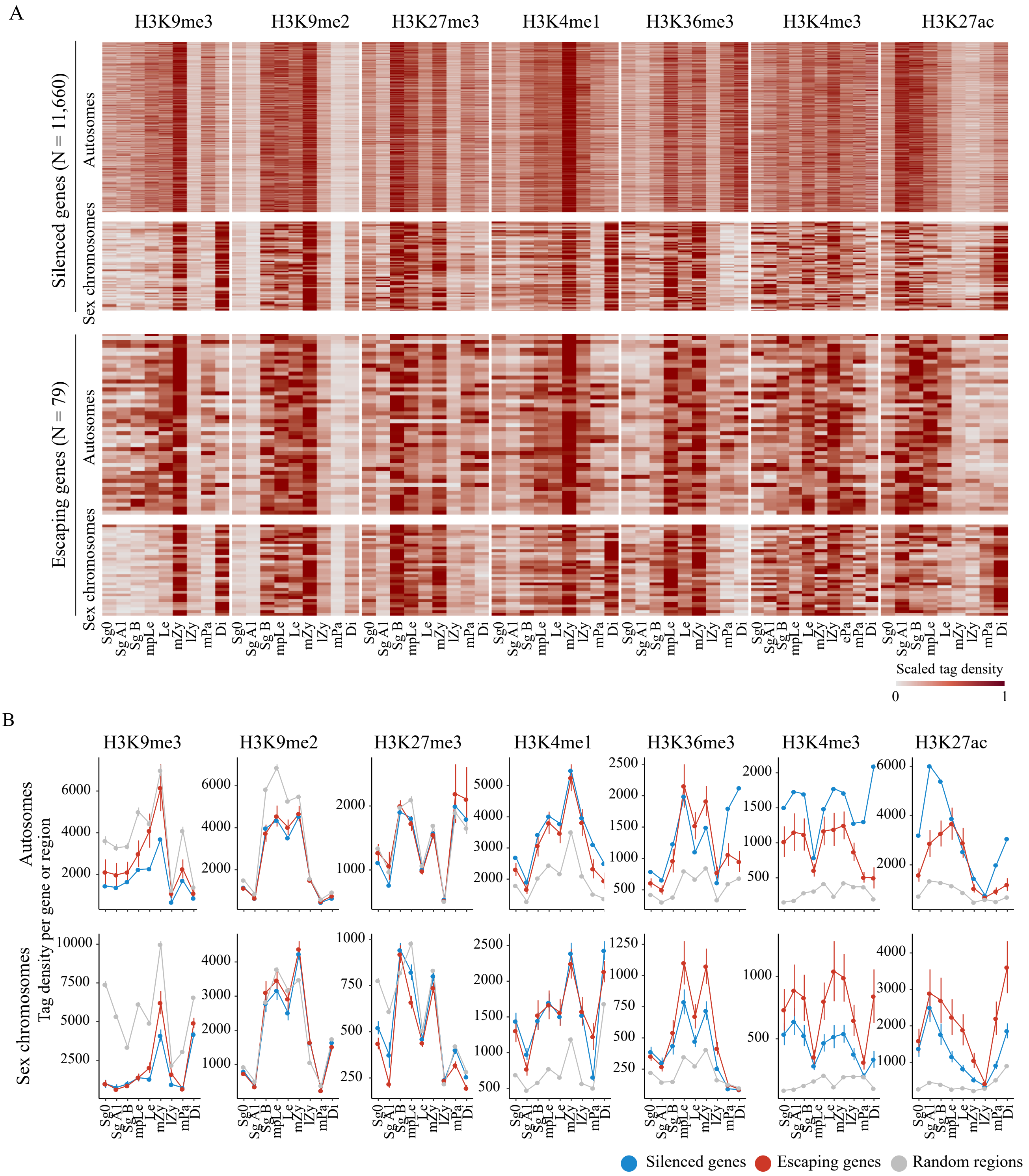
A
H3K9me3
H3K9me2
H3K27me3
H3K4me1
H3K36me3
H3K4me3
H3K27ac
Silenced genes (N = 11,660)
Escaping genes (N = 79)
Autosomes
Sex chromosomes
Sg0
Sg A1
Sg B
mpLe
Le
mZy
lZy
ePa
mPa
Di
Scaled tag density
0
1
B
Autosomes
Sex chromosomes
Tag density per gene or region
Silenced genes
Escaping genes
Random regions

Supp. Fig. S11

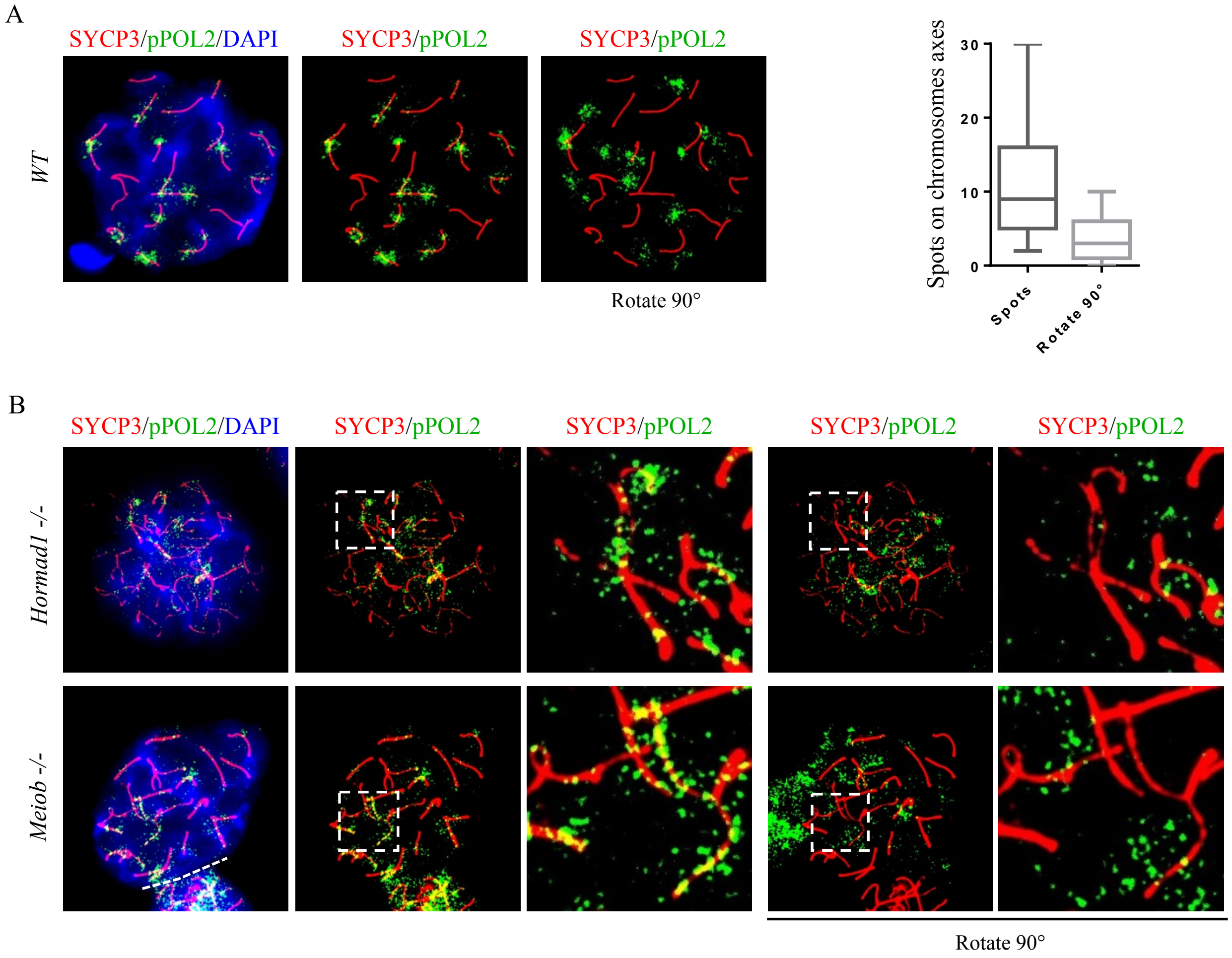
A
SYCP3/pPOL2/DAPI
SYCP3/pPOL2
SYCP3/pPOL2
WT
Rotate 90°
Spots on chromosomes axes
30
20
10
0
Spots
Rotate 90°
B
SYCP3/pPOL2/DAPI
SYCP3/pPOL2
SYCP3/pPOL2
SYCP3/pPOL2
SYCP3/pPOL2
Hormad1 -/-
Meiob -/-
Rotate 90°

Supp. Fig. S12

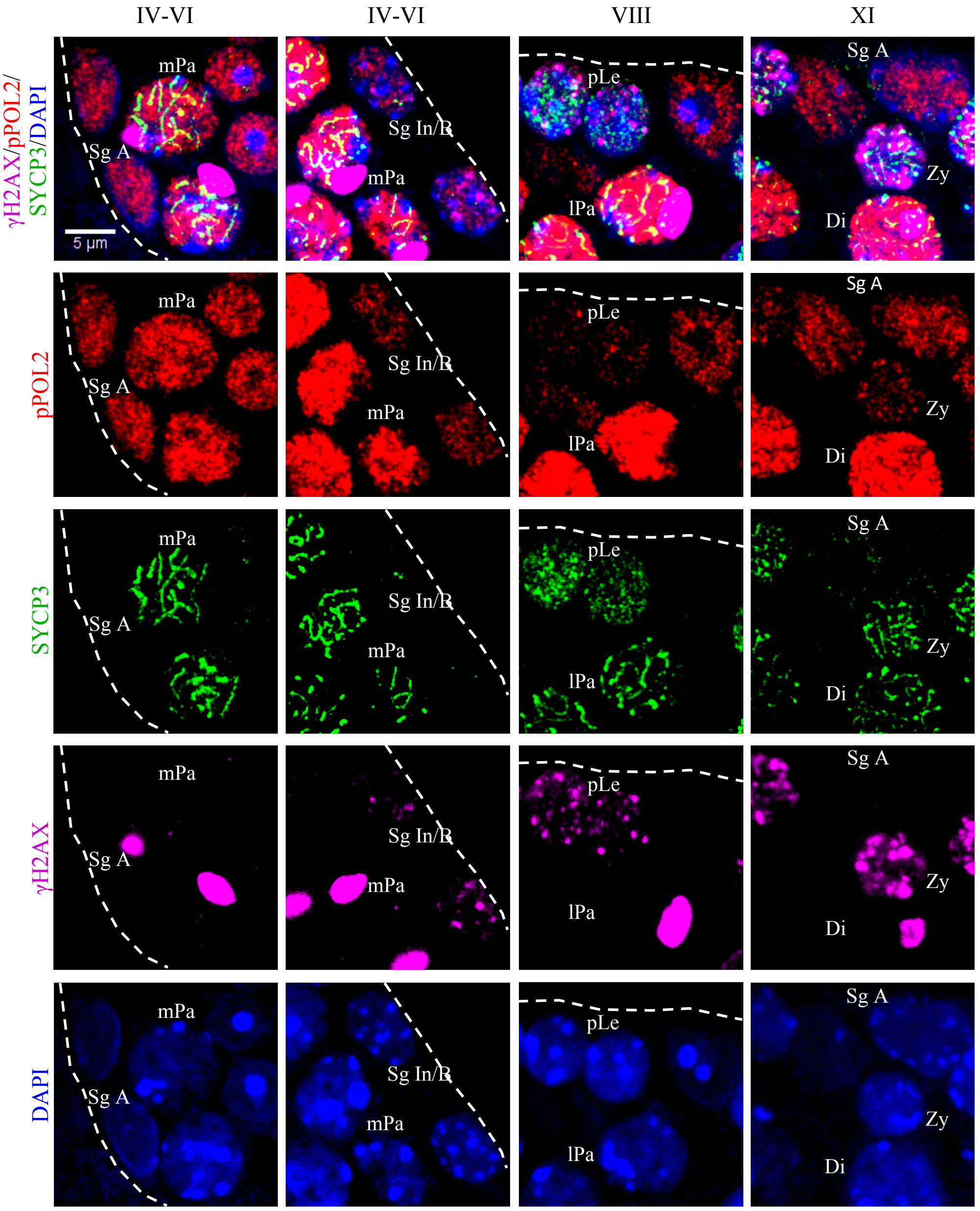

IV-VI
IV-VI
VIII
XI
γH2AX/pPOL2/SYCP3/DAPI
pPOL2
SYCP3
γH2AX
DAPI
5 µm
mPa
Sg A
Sg In/B
pLe
lPa
Zy
Di

Supp. Fig. S13

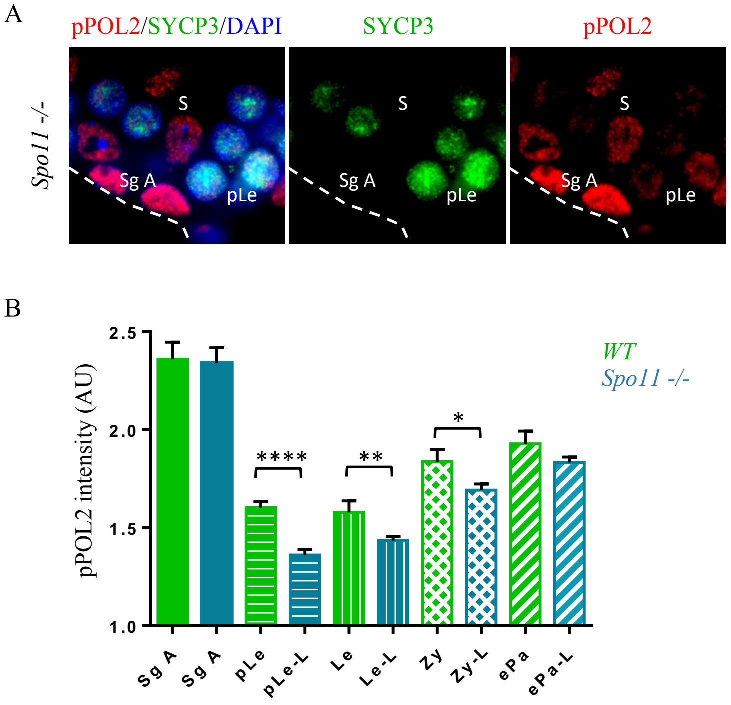
A
pPOL2/SYCP3/DAPI
SYCP3
pPOL2
Spo11 -/-
S
Sg A
pLe
B
pPOL2 intensity (AU)
2.5
2.0
1.5
1.0
WT
Spo11 -/-
****
**
*
Sg A
Sg A
pLe
pLe-L
Le
Le-L
Zy
Zy-L
ePa
ePa-L

Supp. Fig. S14

A

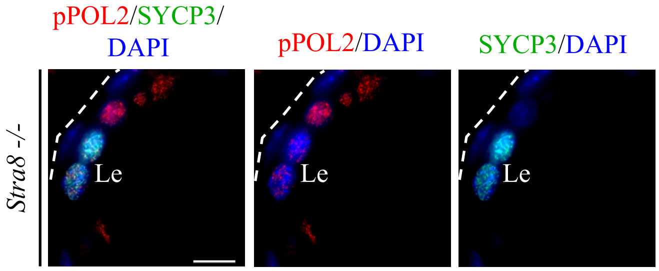

B

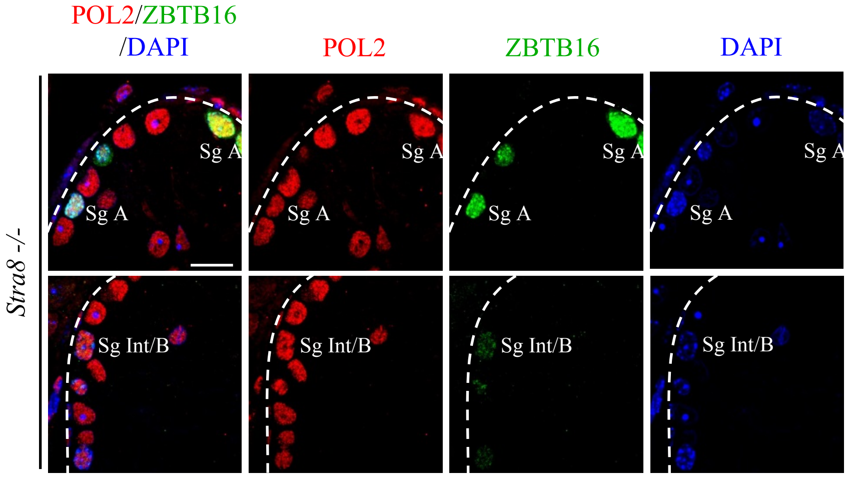

### Supp. Fig. S15

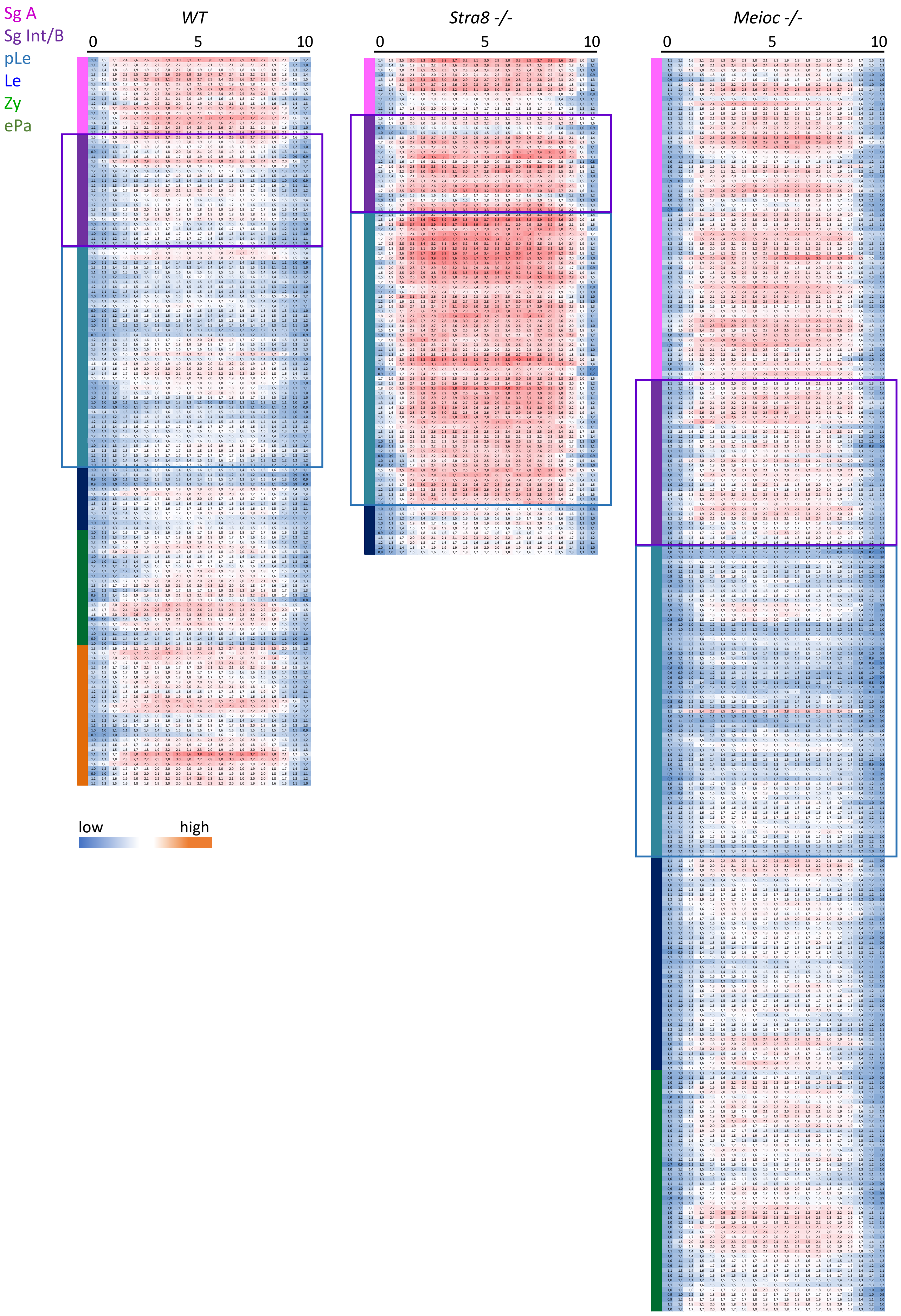

Supp. Fig. S16

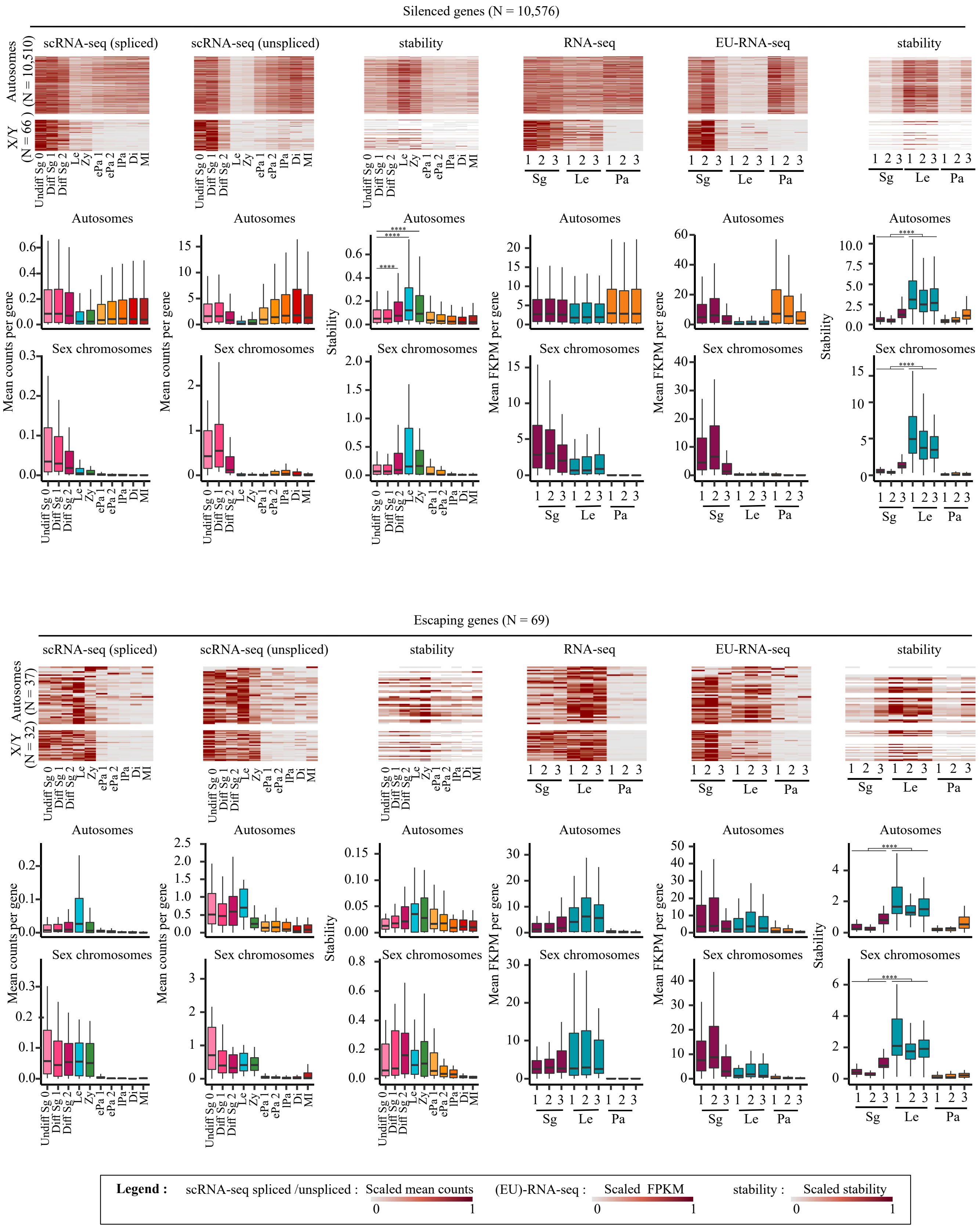
Silenced genes (N = 10,576)
scRNA-seq (spliced)
scRNA-seq (unspliced)
stability
RNA-seq
EU-RNA-seq
stability
Autosomes (N = 10,510)
X/Y (N = 66)
Undiff Sg 0
Diff Sg 1
Diff Sg 2
Le
Zy
ePa 1
ePa 2
lPa
Di
MI
Sg
Le
Pa
Autosomes
Sex chromosomes
Mean counts per gene
Stability
Mean FKPM per gene
****
Escaping genes (N = 69)
Autosomes (N = 37)
X/Y (N = 32)
Legend :
scRNA-seq spliced /unspliced : Scaled mean counts
(EU)-RNA-seq : Scaled FPKM
stability : Scaled stability
0
1

Supp. Fig. S17

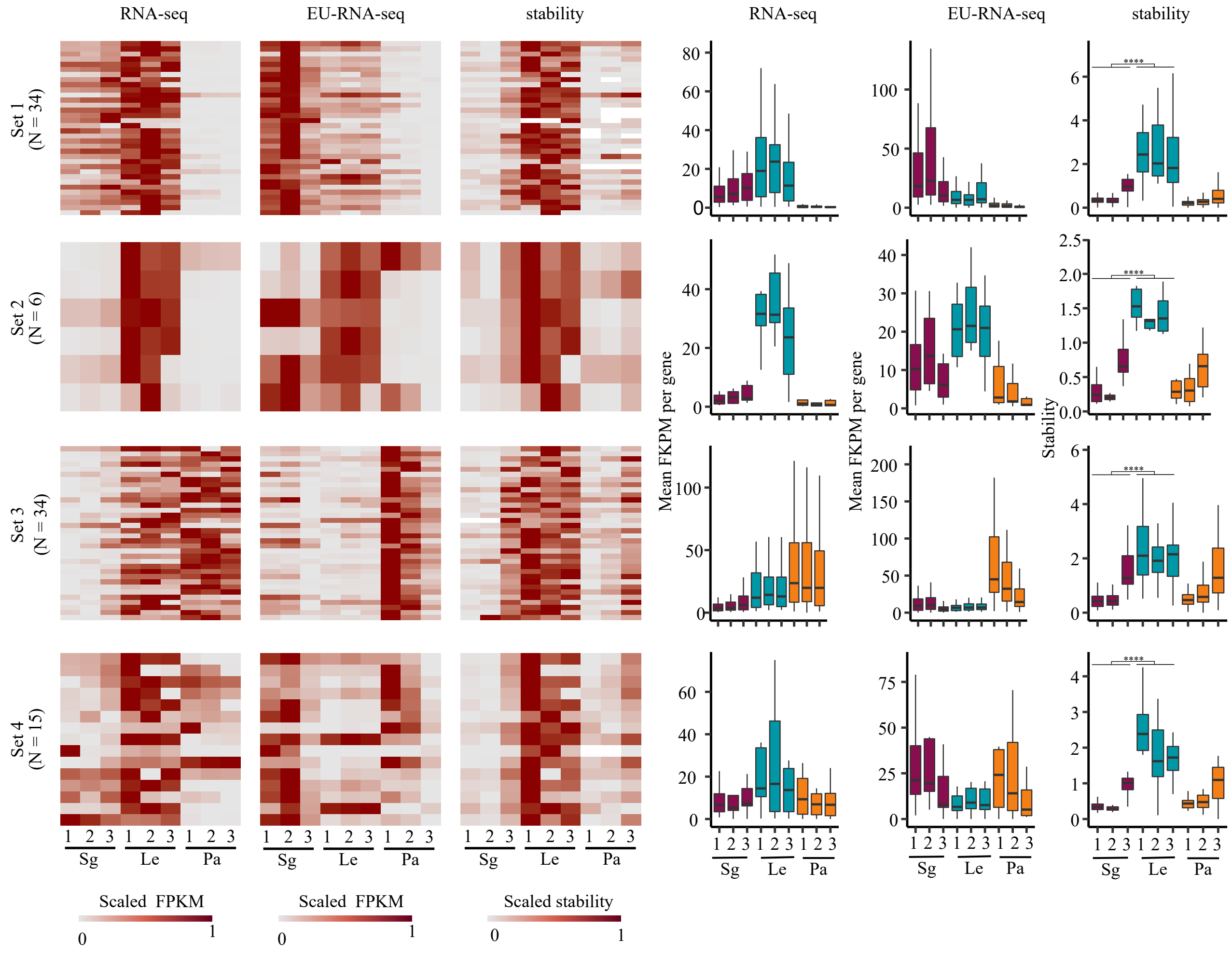

RNA-seq
EU-RNA-seq
stability
Set 1 (N = 34)
Set 2 (N = 6)
Set 3 (N = 34)
Set 4 (N = 15)
1 2 3 1 2 3 1 2 3
Sg Le Pa
Scaled FPKM
Scaled stability
0
1
Mean FKPM per gene
Stability
****
